# Fast and Flexible Estimation of Effective Migration Surfaces

**DOI:** 10.1101/2020.08.07.242214

**Authors:** Joseph H. Marcus, Wooseok Ha, Rina Foygel Barber, John Novembre

## Abstract

An important feature in spatial population genetic data is often “isolation-by-distance,” where genetic differentiation tends to increase as individuals become more geographically distant. Recently, Petkova et al. (2016) developed a statistical method called Estimating Effective Migration Surfaces (EEMS) for visualizing spatially heterogeneous isolation-by-distance on a geographic map. While EEMS is a powerful tool for depicting spatial population structure, it can suffer from slow runtimes. Here we develop a related method called Fast Estimation of Effective Migration Surfaces (FEEMS). FEEMS uses a Gaussian Markov Random Field in a penalized likelihood framework that allows for efficient optimization and output of effective migration surfaces. Further, the efficient optimization facilitates the inference of migration parameters per edge in the graph, rather than per node (as in EEMS). When tested with coalescent simulations, FEEMS accurately recovers effective migration surfaces with complex gene-flow histories, including those with anisotropy. Applications of FEEMS to population genetic data from North American gray wolves shows it to perform comparably to EEMS, but with solutions obtained orders of magnitude faster. Overall, FEEMS expands the ability of users to quickly visualize and interpret spatial structure in their data.

## Introduction

The relationship between geography and genetics has had enduring importance in evolutionary biology (see Felsenstein, 1982). One fundamental consideration is that individuals who live near one another tend to be more genetically similar than those who live far apart (Kimura, 1953, Kimura and Weiss, 1964, Malécot, 1948, Wright, 1943, 1946). This phenomenon is often referred to as “isolation-by-distance” (IBD) and has been shown to be a pervasive feature in spatial population genetic data across many species (Dobzhansky and Wright, 1943, Meirmans, 2012, Slatkin, 1985). Statistical methods that use both measures of genetic variation and geographic coordinates to understand patterns of IBD have been widely applied (Battey et al., 2020, Bradburd and Ralph, 2019). One major challenge in these approaches is that the relationship between geography and genetics can be complex. Particularly, geographic features can influence migration in localized regions leading to spatially heterogeneous patterns of genetic covariation (Bradburd and Ralph, 2019).

Multiple approaches have been introduced to model non-homogeneous IBD in spatial population genetic data (Al-Asadi et al., 2019, Bradburd et al., 2018, Duforet-Frebourg and Blum, 2014, Hanks and Hooten, 2013, McRae, 2006, Petkova et al., 2016, Ringbauer et al., 2018, Safner et al., 2011). Particularly relevant to our proposed approach is the work of Petkova et al. (2016) and Hanks and Hooten (2013). Both approaches model genetic distance using the “resistance distance” on a weighted graph. This distance metric is inspired by concepts of effective resistance in circuit theory models, or alternatively understood as the commute time of a random walk on a weighted graph or as a Gaussian graphical model (specifically a conditional auto-regressive process) (Chandra et al., 1996, Hanks and Hooten, 2013, Rue and Held, 2005). Additionally, the resistance distance approach is a computationally convenient and accurate approximation to spatial coalescent models (McRae, 2006), though it has limitations in asymmetric migration settings (Lundgren and Ralph, 2019).

Hanks and Hooten (2013) introduced a Bayesian model that uses measured ecological covariates, such as elevation, to help predict genetic distances across sub-populations. Specifically, they use a graph-based model for genotypes observed at different spatial locations. Expected genetic distances across sub-populations in their model are given by resistance distances computed from the edge weights. They parameterize the edge weights of the graph to be a function of known biogeographic covariates, linking local geographic features to genetic variation across the landscape.

Concurrently, the Estimating Effective Migration Surfaces (EEMS) method was developed to help interpret and visualize non-homogeneous gene-flow on a geographic map (Petkova et al., 2016, Petkova, 2013). EEMS uses resistance distances to approximate the between-sub-population component of pairwise coalescent times in a “stepping-stone” model of migration and genetic drift (Kimura, 1953, Kimura and Weiss, 1964). EEMS models the within-sub-population component of pairwise coalescent times, with a node-specific parameter. Instead of using known biogeographic covariates to connect geographic features to genetic variation as in Hanks and Hooten (2013), EEMS infers a set of edge weights (and diversity parameters) that explain the genetic distance data. The inference is based on a hierarchical Bayesian model and a Voronoi-tessellation-based prior to encourage piece-wise constant spatial smoothness in the fitted edge weights.

EEMS uses Markov Chain Monte Carlo (MCMC) and outputs a visualization of the posterior mean for effective migration and a measure of genetic diversity for every spatial position of the focal habitat. Regions with relatively low effective migration can be interpreted to have reduced gene-flow over time whereas regions with relatively high migration can be interpreted as having elevated gene-flow. EEMS has been applied to multiple systems to describe spatial genetic structure, but despite EEMS’s advances in computational tractability with respect to the previous work, the MCMC algorithm it uses can be slow to converge, in some cases leading to days of computation time for large datasets (Peter et al., 2018).

These inference problems from spatial population genetics are related to a growing area of interest in the graph signal processing literature referred to as “graph learning” (Dong et al., 2019, Mateos et al., 2019). In graph learning, a noisy signal is measured as a scalar value at a set of nodes from the graph, and the aim is then to infer non-negative edge weights that reflect how spatially “smooth” the signal is with respect to the graph topology (Kalofolias, 2016). In population genetic settings, this scalar could be an allele frequency measured at locations in a discrete spatial habitat with effective migration rates between sub-populations. Like the approach taken by Hanks and Hooten (2013), one widely used representation of smooth graph signals is to associate the smoothness property with a Gaussian graphical model where the precision matrix has the form of a graph Laplacian (Dong et al., 2016, Egilmez et al., 2016). The probabilistic model defined on the graph signal then naturally gives rise to a likelihood for the observed samples, and thus much of the literature in this area focuses on developing specialized algorithms to efficiently solve optimization problems that allow reconstruction of the underlying latent graph. For more information about graph learning and signal processing in general see the excellent survey papers of Dong et al. (2019) and Mateos et al. (2019).

To position the present work in comparison to the “graph learning” literature, our contributions are twofold:

- In population genetics, it is impossible to collect individual genotypes across all the geographic locations and, as a result, we often work with many, often the majority, of nodes having missing data. As far as we are aware, none of the work in graph signal processing considers this scenario and thus their algorithms are not directly applicable to our setting. In addition, if the number of the observed nodes is much smaller than the number of nodes of a graph, one can project the large matrices associated with the graph to the space of observed nodes, therefore allowing for fast and efficient computation.
- On the other hand, highly missing nodes in the observed signals can result in significant degradation of the quality of the reconstructed graph unless it is regularized properly. Motivated by the Voronoi-tessellation-based prior adopted in EEMS (Petkova et al., 2016), we propose regularization that encourages spatial smoothness in the edge weights.

Building on advances in graph learning, we introduce a method, Fast Estimation of Effective Migration Surfaces (FEEMS), that uses optimization rather than MCMC to obtain penalized-likelihood-based estimates of effective migration parameters. In contrast to EEMS which uses a node-specific parameterization of effective migration, we optimize over edge-specific parameters allowing for more flexible migration processes to be fit, such as spatial anisotropy, in which the migration process is not invariant to rotation of the coordinate system (e.g., migration is more extensive along a particular axis). We develop a fast quasi-Newton optimization algorithm (Nocedal and Wright, 2006) and apply it to a dataset of gray wolves from North America. The output is comparable to the results of EEMS but is provided in orders of magnitude less time. With this improvement in speed, FEEMS opens up the ability to perform fast exploratory and iterative data analysis of spatial population structure.

## Results

### Overview of FEEMS

Figure 1 shows a visual schematic of the FEEMS method. The input data are genotypes and spatial locations (e.g., latitudes and longitudes) for a set of individuals sampled across a geographic region. We construct a dense spatial grid embedded in geographic space where nodes represent sub-populations, and we assign individuals to nodes based on spatial proximity (see Supp. Fig. 1 for a visualization of the grid construction and node assignment procedure). The density of the grid is user defined and must be explored to appropriately balance model-mis-specification and computational burden. As the density of the lattice increases, the model is similar to discrete approximations used for continuous spatial processes, but the increased density comes at the cost of computational complexity.

**Figure 1:**
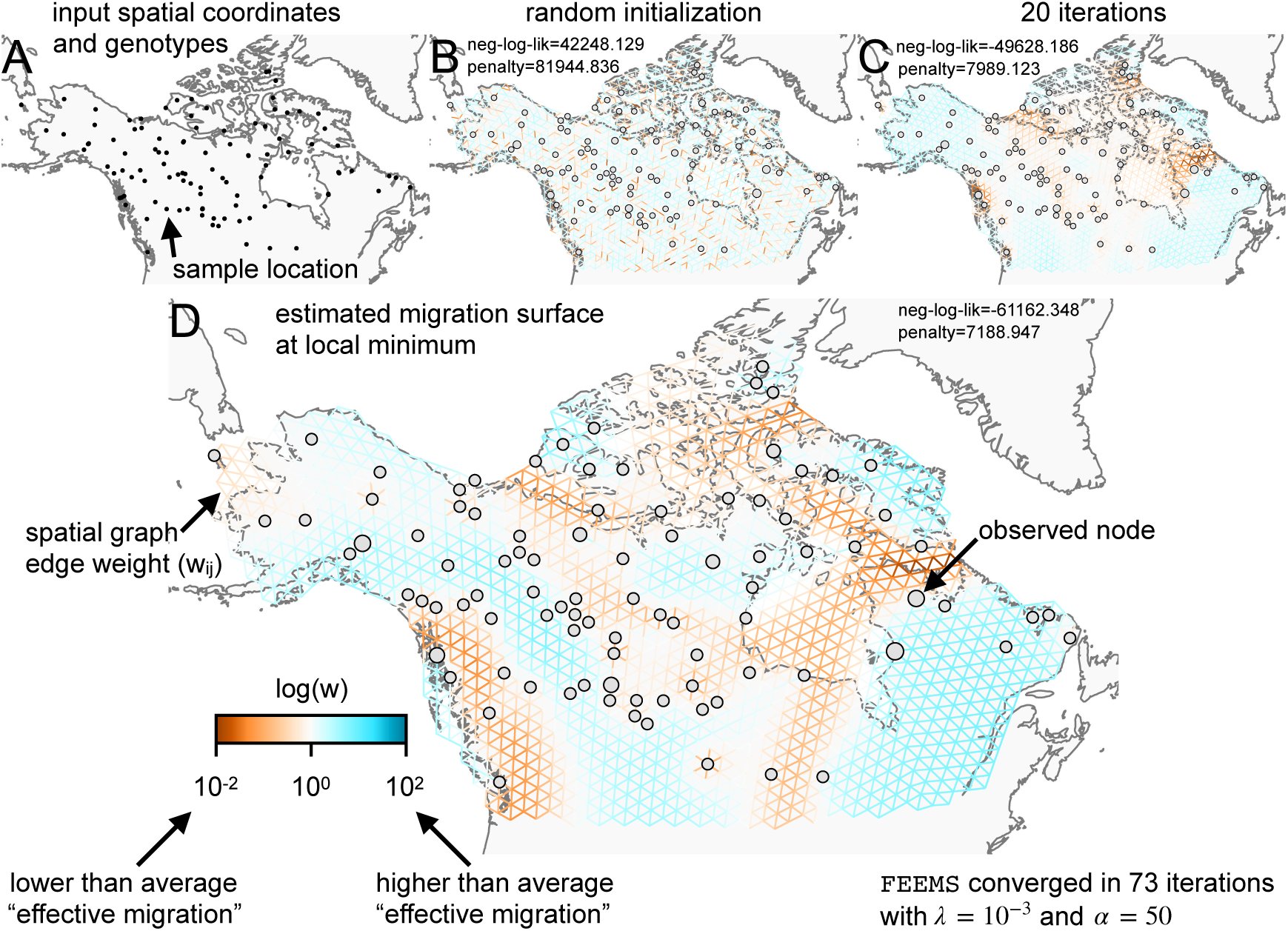
Schematic of the FEEMS model: The full panel shows a schematic of going from the raw data (spatial coordinates and gentoypes) through optimization of the edge weights, representing effective migration, to convergence of FEEMS to a local optima. (A) Map of sample coordinates (black points) from a dataset of gray wolves from North America (Schweizer et al., 2016). The input to FEEMS are latitude and longitude coordinates as well as genotype data for each sample. (B) The spatial graph edge weights after random initialization uniformly over the graph to begin the optimization algorithm. (C) The edge weights after 20 iterations of running FEEMS, when the algorithm has not converged yet. (D) The final output of FEEMS after the algorithm has fully converged. The output is annotated with important features of the visualization.

We assume exchangeability of individuals within each sub-population and estimate allele frequencies, 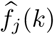, for each sub-population, indexed by *k*, and single nucleotide polymorphism (SNP), indexed by *j*, under a simple Binomial sampling model. We also use the recorded sample sizes at each node to model the precision of the estimated allele frequency. The use of allele frequencies allows a number of advantages in this context: (1) Allele frequencies can be more easily shared between researchers than individual genotypes due to privacy concerns, which is especially relevant in human population genetic studies; (2) We usually gain large computational savings in memory and speed because in most population genetic studies the number of observed locations, in which allele frequencies are estimated, is smaller than the total number of individuals sampled i.e. many individuals are sampled from the same spatial location.

With the estimated allele frequencies in hand, we model the data at each SNP using an approximate Gaussian model whose covariance is shared across all SNPs, in other words we assume that the observed frequencies at each SNP is an independent realization of the same spatial process after rescaling by SNP-specific variation factors. The latent frequency variables, *f*_*j*_ (*k*), are modeled as a Gaussian Markov Random Field (GMRF) with a sparse precision matrix determined by the graph Laplacian and a set of residual variances. The graph’s weighted edges, denoted by *w*_*ij*_ between nodes *i* and *j*, represent gene-flow between the sub-populations (Friedman et al., 2008, Hanks and Hooten, 2013, Petkova et al., 2016). We analytically marginalize out the latent frequency variables and use penalized restricted maximum likelihood to estimate the edge weights of the graph after removing the SNP-specific mean allele frequencies by projecting the data onto contrasts (Felsenstein, 1982, Hanks and Hooten, 2013, Petkova et al., 2016). Our overall goal is to solve the following optimization problem:

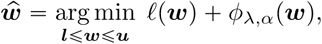

where ***w*** is a vector that stores all the unique elements of the weighted adjacency matrix, ***ℓ*** and ***u*** are element-wise non-negative lower and upper bounds for ***w***, and ℓ(***w***) is the negative log-likelihood function that comes from the GMRF model described above. The penalty, *φ*_*λ,α*_ (***w***), controls how constant or smooth the output migration surface will be and is controlled by the hyperparameters *λ* and *α*. Specifically, the hyperparameters determine a penalty function based on the squared differences between edge weights for pairs of edges that share a common node,

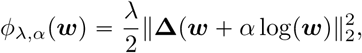

where **Δ** is a signed graph incidence matrix indicating if two edges are connected to the same node. Note that *λ* controls the overall strength of the penalization placed on the output of migration surface while *α* controls the relative strength of the penalization on the logarithmic scale. Thus, if the model is highly penalized, the fitted surface will favor a homogeneous spatial process on the graph across orders of magnitude of edge weights and if the penalty is low, more flexible graphs can be fit, but are potentially prone to over-fitting. Akin to the choice in admixture models of the number of latent ancestral populations or clusters (*K*), inspecting the outputs across a series of *λ* and *α* values is recommended and demonstrated (below). We use sparse linear algebra routines to efficiently compute the objective function and gradient of our parameters, allowing the use of widely applied quasi-Newton optimization algorithms (Nocedal and Wright, 2006) implemented in standard numerical computing libraries like scipy (Virtanen et al., 2020). See the materials and methods section for a detailed description of the statistical models and algorithms used.

### Evaluating FEEMS on “out of model” coalescent simulations

While our statistical model is not directly based on a population genetic process, it is useful to see how it performs on simulated data under the coalescent stepping stone model. In these simulations we know, by construction, the model we fit (FEEMS) is different from the true model we simulate data under (the coalescent), allowing us to assess the robustness of the fit to a controlled form of model mis-specification. In Figure 2 we use `msprime` (Kelleher et al., 2016) to recapitulate and extend the results of Petkova et al. (2016), simulating data under the coalescent in three simple migration scenarios with two different spatial sampling designs. Note that in Supp. Fig. 2 we display a larger set of simulations with additional sampling configurations. For brevity, here we only show results for *λ =*.001 and *α =* 50, based on values that performed well after experimental tuning. In Supp. Fig. 3 and Supp. Fig. 4, we also show results varying *λ* and *α* for two migration scenarios with one particular sampling design.

**Figure 2:**
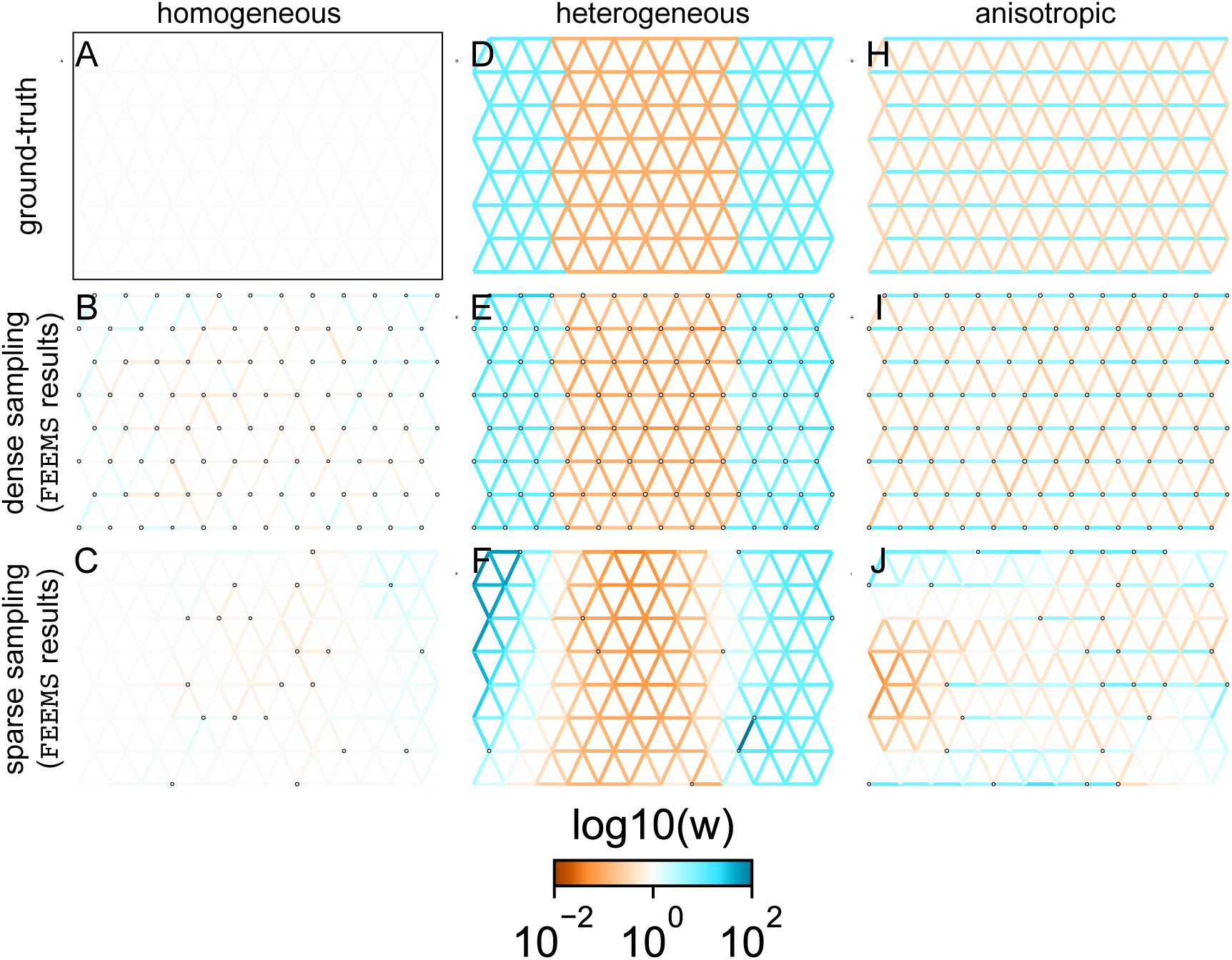
FEEMS fit to coalescent simulations: We run FEEMS on coalescent simulations, varying the migration history (columns) and sampling design (rows). The first column (A-C) shows the ground-truth and fit of FEEMS to coalescent simulations with a homogeneous migration history i.e. a single migration parameter for all edge weights. Note that the ground-truth simulation figures (A,D,F) display coalescent migration rates, not fitted effective migration rates output by FEEMS. The second column (D-F) shows the ground truth and fit of FEEMS to simulations with a heterogeneous migration history i.e. reduced gene-flow, with 10 fold lower migration, in the center of the habitat. The third column (H-J) shows the ground truth and fit of FEEMS to an anisotropic migration history with edge weights facing east-west having five fold higher migration than north-south. The second row (B,E,H) shows a sampling design with no missing observations on the graph. The final row (C,F,I) shows a sampling design with 80% of nodes missing at random.

**Figure 3:**
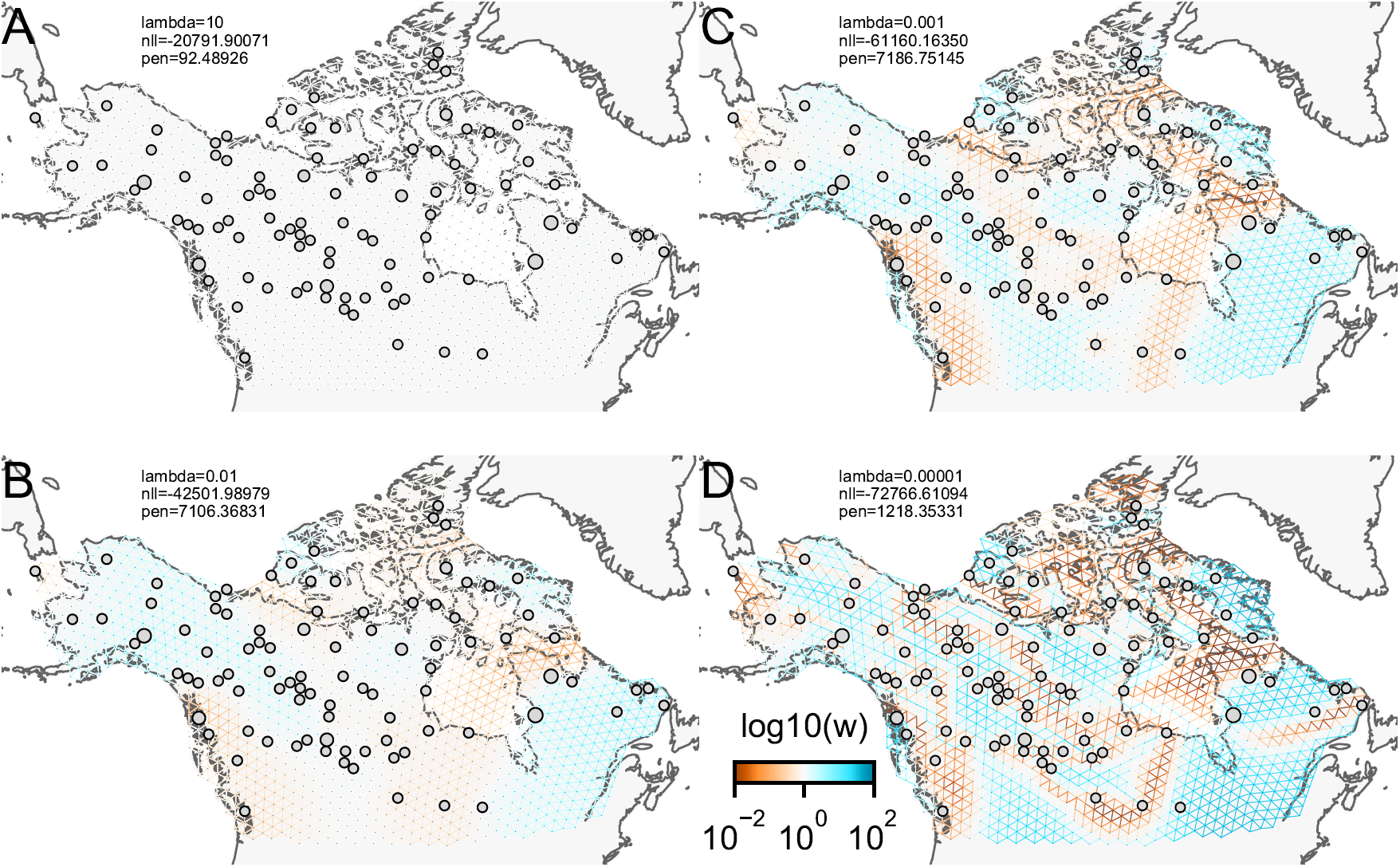
The fit of FEEMS to the North American gray wolf dataset for different choices of the smoothing regularization parameter *λ*: (A) *λ =* 10, (B) *λ =* 10^−2^, (C) *λ =* 10^−3^, and (D) *λ =* 10^−5^. As expected, when *λ* decreases from large to small (A-D), the fitted graph becomes less smooth and presumably eventually over-fits to the data, revealing a patchy surface in (D), whereas, earlier in the regularization path FEEMS fits a completely homogeneous surface with all edge weights having the same fitted value, like in (A).

**Figure 4:**
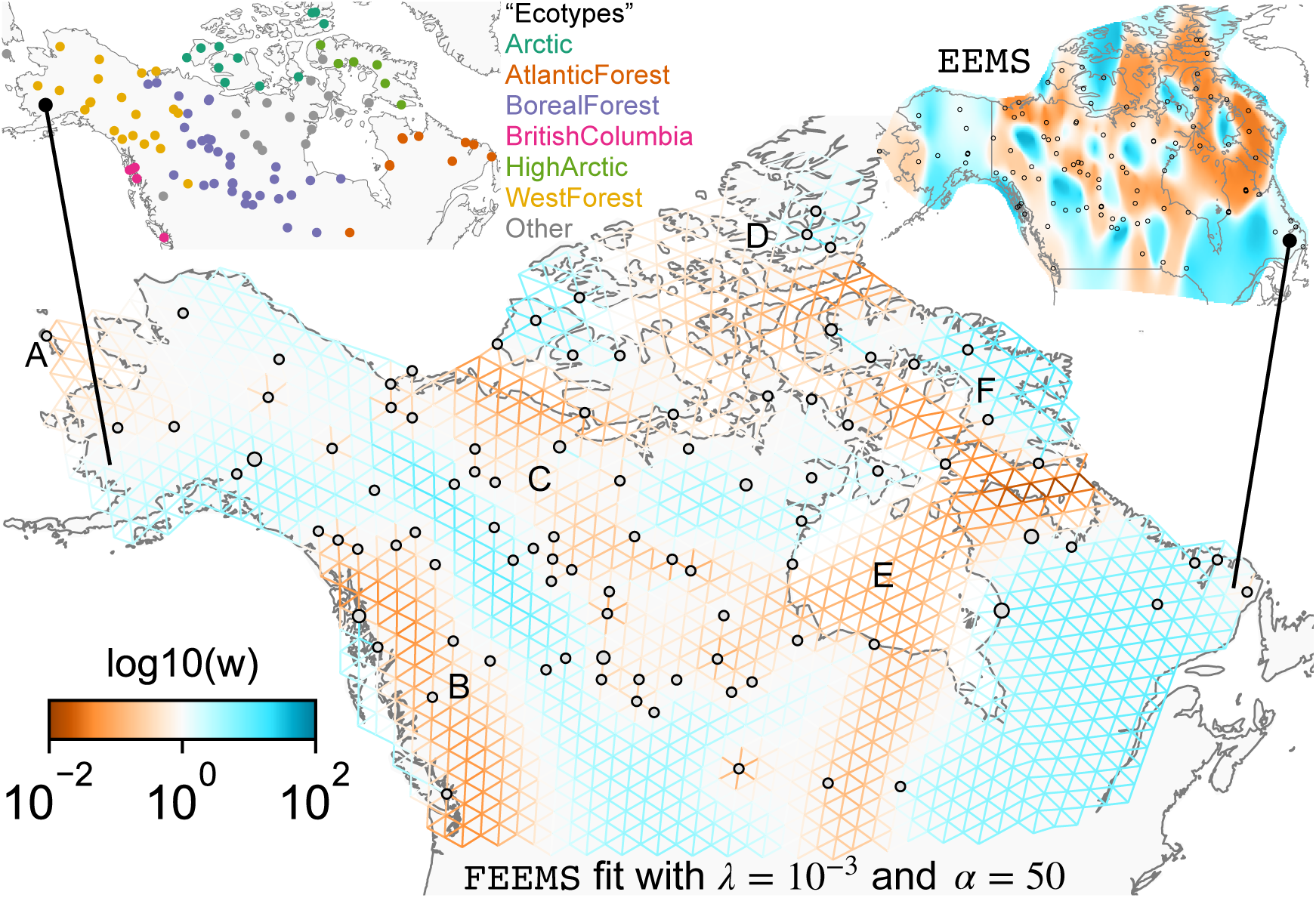
FEEMS applied to a population genetic dataset of North American gray wolves: We show the fit of FEEMS to a previously published dataset of North American gray wolves. This fit corresponds to a setting of tuning parameters at *λ =* 10^−3^, *α =* 50. We show the fitted parameters in log-scale with lower effective migration shown in orange and higher effective migration shown in blue. The bold text letters highlights a number of known geographic features that could have plausibly influenced Wolf migration over time: (A) St. Lawerence Island, (B) Coastal mountain ranges in British Columbia, (C) The boundary of Boreal Forest and Tundra eco-regions in the Shield Taiga, (D) Queen Elizabeth Islands, (E) Hudson Bay, and (F) Baffin Island. We also display two insets to help interpret the results and compare them to EEMS. In the top left inset we show a map of sample coordinates colored by an ecotype label provided by Schweizer et al. (2016). These labels were devised using a combination of genetic and ecological information for 94 “un-admixed” gray wolf samples, and the remaining samples were labeled “Other”. We can see these ecotype labels align well with the visualization output provided by FEEMS. In the right inset we display a visualization of the posterior mean effective migration rates from EEMS.

The first migration scenario (Figure 2A-C) is a spatially homogeneous model where all the migration rates are set to be a constant value on the graph, this is equivalent to simulating data under an homogeneous isolation-by-distance model. In the second migration scenario (Figure 2D-E) we simulate a non-homogeneous process by representing a geographic barrier to migration, lowering the migration rates by a factor of 10 in the center of the habitat relative to the left and right regions of the graph. Finally, in the third migration scenario (Figure 2G-I) we simulate a pattern which corresponds to anisotropic migration with edges that point east/west being assigned to a five-fold higher migration rate than edges pointing north/south. For each migration scenario we simulate two sampling designs. In the first “dense-sampling” sampling design (Figure 2B,E,I) we sample individuals for every node of the graph. Next, in the “sparse-samplng” sampling design (Figure 2C,F,J) we randomly sample individuals for only 20% of the nodes.

As expected, FEEMS performs best when all the nodes are sampled on the graph, i.e. when there is no missing data (Figure 2B,E,H). Interestingly, in the simulated scenarios with many missing nodes, FEEMS can still partly recover the migration history, including the presence of anisotropic migration (Figure 2F). A sampling scheme with a central gap leads to a slightly narrower barrier in the heterogeneous migration scenario (Supp. Fig. 2I) and for the anisotropic scenario, a degree of over-smoothness in the northern and southern regions of the center of the graph (Supp. Fig. 2N). For the missing at random sampling design, FEEMS is able to recover the relative edge weights surprisingly well for all scenarios, with the inference being the most challenging when there is anisotropic migration. We emphasize that the potential for FEEMS to recover anisotropic migration is novel relative to EEMS, which was parameterized for fitting non-stationary isotropic migration histories and produces banding patterns perpendicular to the axis of migration when applied to data from anisotropic coalescent simulations (see Petkova et al. (2016) supplementary figure 2; see also Supp. Note “*Edge versus node parameterization*” for a related discussion). Overall, even with sparsely sampled graphs, FEEMS is able to produce visualizations that qualitatively capture the migration history in “out of model” coalescent simulations.

### Application of FEEMS to genotype data from North American gray wolves

To assess the performance of FEEMS on real data we used a previously published dataset of 111 gray wolves sampled across North America typed at 17, 729 SNPs (Schweizer et al., 2016), Supp. Fig. 5). This dataset has a number of advantageous features that make it a useful test case for evaluating FEEMS: (1) The broad sampling range across North America includes a number of relevant geographic features that, a priori, could conceivably lead to restricted gene-flow averaged throughout the population history. These geographic features include mountain ranges, lakes and island chains. (2) The scale of the data is consistent with many studies for non-model systems whose spatial population structure is of interest. For instance, the relatively sparse sampling leads to a challenging statistical problem where there is the potential for many unobserved nodes (sub-populations), depending the density of the grid chosen. Before applying FEEMS, we confirmed a signature of spatial structure in the data through regressing genetic distances on geographic distances and top genetic PCs against geographic coordinates (Supp. Fig. 6, 7, 8, 9).

We ran FEEMS with four different values of the smoothness parameter, *λ* (from large *λ =* 10 to small *λ =* 10^−5^, while setting the tuning parameter *α* to a value that we found that worked for multiple data applications and simulations (*α =* 50, Figure 3). One interpretation of our regularization penalty is that it encourages fitting models of homogeneous and isotropic migration. When *λ* is very large (Figure 3A), we see FEEMS fits a model where all of the edge weights on the graph nearly equal the mean value, hence all the edge weights are colored white in the relative log-scale. In this case, FEEMS is fitting a completely homogeneous migration model where all the estimated edge weights get assigned the same value on the graph. Next, as we sequentially lower the penalty parameter and (Figure 3B,C,D) the fitted graph begins to appear more complex and heterogeneous as expected (discussed further below).

We also ran multiple replicates of ADMIXTURE for *K =* 2 to *K =* 8, selecting for each *K* the highest likelihood run among replicates to visualize (Supp. Fig. 10). As expected in a spatial genetic dataset, nearby samples have similar admixture proportions and continuous gradients of changing ancestries are spread throughout the map (Bradburd et al., 2018). Whether such gradients in admixture coefficients are due to isolation by distance or specific geographic features that enhance or diminish the levels of genetic differentiation is an interpretive challenge. Explicitly modeling the spatial locations and genetic distance jointly using a method like EEMS or FEEMS is exactly designed to explore and visualize these types of questions in the data (Petkova et al., 2016, Petkova, 2013).

Once we have run FEEMS for a grid of regularization parameters it is helpful to look more closely at particular solutions that find a balance between spatial homogeneity and complexity (Figure 4). Spatial features in the FEEMS visualization qualitatively matches the structure plot output from ADMIXTURE using *K =* 6 (Supp. Fig. 10). We add labels on the figure to highlight a number of pertinent features: (A) St. Lawrence Island, (B) the coastal islands and mountain ranges in British Columbia, (C) The boundary of Boreal Forest and Tundra eco-regions in the Shield Taiga, (D) Queen Elizabeth Islands, (E) Hudson Bay, and (F) Baffin Island. Many of these features were described in Schweizer et al. (2016) by interpretation of ADMIXTURE, PCA, and *F*_*ST*_ statistics. FEEMS is able to succinctly provide an interpretable view of these data in a single visualization. Indeed many of these geographic features plausibly impact gray wolf dispersal and population history (Schweizer et al., 2016).

### Comparison to EEMS

We also ran EEMS on the same gray wolf dataset described throughout this manuscript. We used default parameters provided by EEMS but set the number of burn-in iterations to 20 &3×00D7; 10^6^, MCMC iterations to 50 × 10^6^, and thinning intervals to 2000. We were unable to run EEMS in a reasonable run time (⩽3 days) for the dense spatial grid of 1207 nodes so we ran EEMS and FEEMS on a sparser graph with 307 nodes.

We find that FEEMS is multiple orders of magnitude faster than EEMS, even when running multiple runs of FEEMS for different regularization settings on both the sparse and dense graphs (Table 1). The total FEEMS run-times in Table 1 also include the time needed to construct relevant graph data structures and initialization. We note that constructing the graph and fitting the model with very low regularization parameters are the most computationally demanding steps in running FEEMS.

**Table 1:** Runtimes for FEEMS and EEMS on the North American gray wolf dataset: We show a table of runtimes for FEEMS and EEMS for two different grid densities, a sparse grid with 307 nodes and a dense grid with 1207 nodes. In the first two rows we show the total runtimes for both EEMS and FEEMS. In the following rows we show the total runtime for FEEMS, broken down into multiple components i.e. initialization time and the time to fit four solutions with different smoothing parameters.

| Method | Sparse Grid (run-time) | Dense Grid (run-time) |
| --- | --- | --- |
| EEMS | 27.43hrs | N/A |
| FEEMS (total) | 13.02s | 3.54min |
| FEEMS (init) | 8.25s | 2min 11s |
| FEEMS ($\lambda = 10$) | 604ms | 10.7s |
| FEEMS ($\lambda = 10^{-2}$) | 442ms | 7.78s |
| FEEMS ($\lambda = 10^{-3}$) | 917ms | 9.18s |
| FEEMS ($\lambda = 10^{-5}$) | 2.81s | 53.9s |

We find that many of the same geographic features that have reduced or enhanced gene-flow are concordant between the two methods. The EEMS visualization, qualitatively, best matches solutions of FEEMS with lower regularization penalties (Figure 4, Supp. Fig. 11); however, based on the ADMIXTURE results and visual inspection in relation to known geographical features, we find these solutions to be less satisfying compared to those with higher penalties and believe the solutions output from lower penalties are likely overfitting the data. Indeed, we only see a small gain in the *R*_2_ when comparing observed and fitted distances computed from the output graphs of Figure 3C and Figure 3D (Supp. Fig. 6). We note that in many of the EEMS runs the MCMC appears to not have converged (based on visual inspection of trace plots) even after a large number of iterations.

## Discussion

FEEMS is a fast approach that provides an interpretable view of spatial population structure in real datasets and simulations. We want to emphasize that beyond being a fast optimization approach for inferring population structure, our parameterization of the likelihood opens up a number of exciting new directions for improving spatial population genetic inference. Notably, one major difference between EEMS and FEEMS is that in FEEMS each edge weight is assigned its own parameter to be estimated whereas, in EEMS, each node is assigned a parameter and each edge is constrained to be the average effective migration between the nodes it connects (see Materials and Methods and Supp. Note “*Edge versus node parameterization*” for details). The node-based parameterization in EEMS makes it difficult to incorporate anisotropy and asymmeteric migration (Lundgren and Ralph, 2019). As we have shown here, FEEMS’s simple and novel parameterization already has potential to fit anisotropic migration (as shown in coalescent simulations) and may be extendable to other more complex migration processes (such as long-range migration, see below).

FEEMS estimates one set of graph edge weights for each setting of the tuning parameters *λ* and *α* which control the smoothness of the fitted edge-weights. One general challenge, which is not unique to this method, is selecting a particular set of tuning parameters. A natural approach is to use cross-validation, which estimates the out-of-sample fit of FEEMS for a particular model (selection of *λ* and *α*). While cross-validation might be useful for assessing the choice of tuning parameters, in preliminary experiments applying cross validation by holding out individuals or observed nodes, and assessing performance via the model-likelihood, we found too much variation across cross-validation folds to reliably tune *λ* and *α* (results not shown). In order to reduce the variation across different folds, we also applied cross-validation with standardization (Bradburd et al., 2018), where the model-likelihood is standardized for each fold, and approximate leave-one-out cross-validation Wilson et al. (2020), where the leave-one-out CV likelihood is approximated with a few steps of the quasi-Newton algorithm warm-started from the full training set migration surfaces. Neither of these approaches were promising for reliable model selection. We suspect this poor performance is due to spatial dependency of allele frequencies and the large fraction of unobserved nodes. In unsupervised learning settings like this one, it is not obvious that estimates of out of sample fit will always lead to the most biologically interpretable models and sometimes other metrics can be preferable, such as those based on the stability of the solution to perturbations like variable initialization (Wu et al., 2016). Stability-based approaches for model selection could be a fruitful future direction to develop a formal procedure for tuning. Currently, we recommend fitting FEEMS with several values of the tuning parameters and interpreting the results in an integrative fashion with other analyses.

We find it useful to fit FEEMS to a sequential grid of regularization parameters and to look at what features are consistent and vary across multiple fits. Informally, one can gain an indication of the strongest features in the data by looking at the order they appear in the regularization path i.e. what features overcome the strong penalization of smoothness in the data and that are highly supported by the likelihood. For example, early in the regularization path, we see regions of reduced gene-flow occurring in the west coast of Canada that presumably correspond to Coastal mountain ranges and islands in British Columbia (Figure 3B) and this reduced gene-flow appears throughout more flexible fits with lower *λ*.

Beyond tuning the unknown parameters, we encountered other challenges when solving this difficult optimization problem. Notably, the objective function we optimize is non-convex so any visualization output by FEEMS should be considered a local optimum and, as a result, with different initialization we could get different results. Overall, we found the output visualization was not sensitive to initialization, and thus our default setting is constant initialization fitted under an homogeneous isolation by distance model (See Materials and Methods).

When comparing to EEMS, we found FEEMS to be much faster (Table 1). While this is encouraging, care must be taken because the goals and outputs of FEEMS and EEMS have a number of differences. FEEMS fits a sequential grid of solutions for different regularization parameters whereas EEMS infers a posterior distribution and outputs the posterior mean as a point estimate. So in order to compare the results, in principal, one must compare many FEEMS visualizations to a single EEMS visualization. FEEMS is not a Bayesian method and unlike EEMS, which explores the entire landscape of the posterior distribution, FEEMS returns a particular point estimate: a local minimum point of the optimization landscape. Setting the prior hyper-parameters in EEMS act somewhat like a choice of tuning parameters, except that EEMS uses hierarchical priors that in principle allow for exploration of multiple scales of spatial structure in a single run; this arguably results in less sensitivity to user-based settings but requires potentially long computation times for adequate MCMC convergence.

One natural extension to FEEMS, pertinent to a number of biological systems, is incorporating long-range migration (Bradburd et al., 2016, Pickrell and Pritchard, 2012). In this work we have used a triangular lattice embedded in geographic space and enforced smoothness in nearby edge weights through penalizing their squared differences (see Materials and Methods). We could imagine changing the structure of the graph by adding edges to allow for long-range connection; however our current regularization scheme would not be appropriate for this setting. Instead, we could imagine adding an additional penalty to the objective, which would only allow a few long range connections to be tolerated. This could be considered to be a combination of two existing approaches for graph-based inference, graphical lasso (GLASSO) and graph Laplacian smoothing, combining the smoothness assumption for nearby connections and the sparsity assumption for long-range connections (Friedman et al., 2008, Wang et al., 2016). Another potential methodological avenue to incorporate long-range migration is to use a “greedy” approach. We could imagine adding long-range edges one a time, guided by re-fitting the spatial model and taking a data driven approach to select particular long-range edges to include. The proposed greedy approach could be considered to be a spatial graph analog of TreeMix (Pickrell and Pritchard, 2012).

Another interesting extension would be to incorporate asymmetric migration into the framework of resistance distance and Gaussian Markov Random Field based models. Recently, Hanks (2015) developed a promising new framework for deriving the stationary distribution of a continuous time stochastic process with asymmetric migration on a spatial graph. Interestingly, the expected distance of this process has a similar “flavor” to the resistance distance based models, in that it depends on the pseudo-inverse of a function of the graph Laplacian. Hanks (2015) used MCMC to estimate the effect of known covariates on the edge weights of the spatial graph. Future work could adapt this framework into the penalized optimization approach we have considered here, where adjacent edge weights are encouraged to be smooth.

Finally, when interpreted as mechanistic rather than statistical models, both EEMS and FEEMS implicitly assume time-stationarity, so the estimated migration parameters should be considered to be “effective” in the sense of being averaged over time in a reality where migration rates are dynamic and changing (Pickrell and Reich, 2014). The MAPS method is one recent advance that utilizes long stretches of shared haplotypes between pairs of individuals to perform Bayesian inference of time varying migration rates and population sizes (Al-Asadi et al., 2019). With the growing ability to extract high quality DNA from ancient samples, another exciting future direction would be to apply FEEMS to ancient DNA datasets over different time transects in the same focal geographic region to elucidate changing migration histories (Mathieson et al., 2018). There are a number of technical challenges in ancient DNA data that make this a difficult problem, particularly high levels of missing and low-coverage data. Our modeling approach could be potentially more robust, in that it takes allele frequencies as input, which may be estimable from dozens of ancient samples at the same spatial location, in spite of high degrees of missingness (Korneliussen et al., 2014).

In closing, we look back to a review titled “How Can We Infer Geography and History from Gene Frequencies?” published in 1982 (Felsenstein, 1982). In this review, Felsenstein laid out fundamental open problems in statistical inference in population genetic data, a few of which we restate as they are particularly motivating for our work:

- “For any given covariance matrix, is there a corresponding migration matrix which would be expected to lead to it? If so, how can we find it?”
- “How can we characterize the set of possible migration matrices which are compatible with a given set of observed covariances?”
- “How can we confine our attention to migration patterns which are consistent with the known geometric co-ordinates of the populations?”
- “How can we make valid statistical estimates of parameters of stepping stone models?”

The methods developed here aim to help address these longstanding problems in statistical population genetics and to provide a foundation for future work to elucidate the role of geography and dispersal in ecological and evolutionary processes.

## Materials and Methods

### Model description

See Supp. Note “*Mathematical notation*” for a detailed description of the notation used to describe the model. To visualize and model spatial patterns in a given population genetic dataset, FEEMS uses an undirected graph, *𝒢 =* (𝒱, ℰ) with |*V*| = *d*, where nodes represent sub-populations and edge weights (*w*_*ij*_) _*(i,j*)∈*ℰ*_ represent the level of gene-flow between sub-populations *i* and *j*. For computational convenience, we assume *𝒢* is a highly sparse graph, specifically a triangular grid that is embedded in geographic space around the sample coordinates. We observe a genotype matrix, ***Y ∈*** *R*^*n×p*^, with *n* rows representing individuals and *p* columns representing SNPs. We imagine diploid individuals are sampled on the nodes of *𝒢* so that *y*_*ij*_ (*k*) ∈ {0, 1, 2} records the count of some arbitrarily predefined allele in individual *i*, SNP *j*, on node *k ∈* 𝒱. We assume a commonly used simple Binomial sampling model for the genotypes:

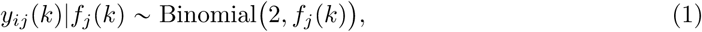

where conditional on *f*_*j*_ (*k*) for all *j, k*, the *y*_*ij*_ (*k*) ‘s are independent. We then estimate an allele frequency at each node and SNP by maximum likelihood:

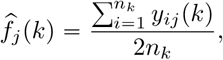

where *n*_*k*_ is the number of individuals sampled at node *k*. We estimate allele frequencies at *o* of the observed nodes out of *d* total nodes on the graph. From (1), the estimated frequency in a particular sub-population, conditional on the latent allele frequency, will approximately follow a Gaussian distribution:

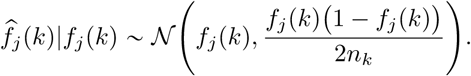

Using vector notation, we represent the joint model of estimated allele frequencies as:

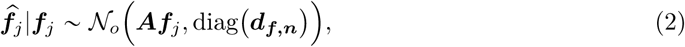

where 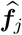 is a *o ×* 1 vector of estimated allele frequencies at observed nodes, ***f***_*j*_ is a *d* × 1 vector of latent allele frequencies at all the nodes (both observed and unobserved), and ***A*** is a *o* ×*d* node assignment matrix where ***A***_*kℓ*_ *=* 1 if the *k*th estimated allele frequency comes from sub-population ℓ and ***A***_*kℓ*_ 0 otherwise; and diag (***d***_***f***,***n***_ ) denotes a *o × o* diagonal matrix whose diagonal elements corresponds to the appropriate variance term at observed nodes.

To summarize, we estimate allele frequencies from a subset of nodes on the graph and define latent allele frequencies for all the nodes of the graph. The assignment matrix ***A*** maps these latent allele frequencies to our observations. Our summary statistics (the data) are thus 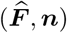 where 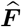 is a *o × p* matrix of estimated allele frequencies and ***n*** is a *o ×*1 vector of sample sizes for every observed node. We assume the latent allele frequencies come from a Gaussian Markov Random Field:

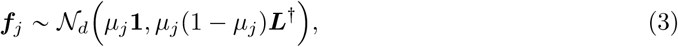

where ***L*** is the graph Laplacian and *µ*_*j*_ represents the average allele frequency across all of the sub-populations. Note that the multiplication by the SNP-specific factor *µ*_*j*_ (1 − *µ*_*j*_) ensures that the variance of the latent allele frequencies vanishes as the average allele frequency approaches to 0 or 1. One interpretation of this model is that the expected squared Euclidean distance between latent allele frequencies on the graph, after being re-scaled by (*µ*_*j*_ 1 −*µ*_*j*_), is exactly the resistance distance of an electrical circuit (Hanks and Hooten, 2013, McRae, 2006):

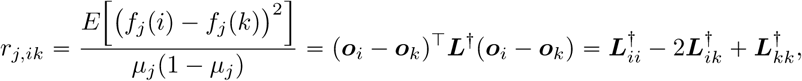

where ***o***_*i*_ is a one-hot vector (i.e., storing a 1 in element *i* and zeros elsewhere). It is known that the resistance distance is equivalent to the expected commute time between nodes *i* and *k* of a random walker on the weighted graph *𝒢* (Chandra et al., 1996). Additionally, the model (3) forms a Markov random field, and thus any latent allele frequency *f*_*j*_ *(i)* is conditionally independent of all other allele frequencies given its neighbors which are encoded by nonzero elements of ***L*** (Koller and Friedman, 2009, Lauritzen, 1996).^1^

Using the law of total variance formula, we can derive from (2), (3) an analytic form for the marginal likelihood. Before proceeding, however, we further approximate the model by assuming 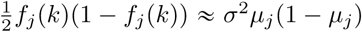 for all *j* and *k*. This assumption is mainly for computational purposes and may be a coarse approximation in general. On the other hand, the assumption is not too strong if we exclude SNPs with extremely rare allele frequencies, and more importantly, we find it leads to a good empirical performance, both statistically and computationally. With this approximation the residual variance parameter *σ*^2^ is still unknown and needs to be estimated.

With the above considerations, we arrive at the following marginal likelihood:^2^

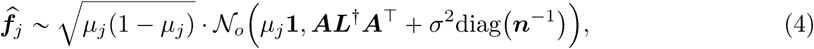

where diag ***n***^−1^ is a *o × o* diagonal matrix computed from the sample sizes at observed nodes. To remove the SNP means we transform the estimated frequencies by a contrast matrix, ***C ∈****R*^(p*−1)×o*^, that is orthogonal to the one-vector:

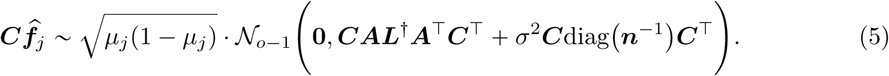

Letting 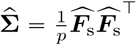 be the *o ×o* sample covariance matrix of estimated allele frequencies after rescaling, i.e. 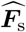 is a matrix formed by rescaling the columns of 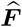 by 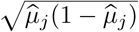, where 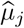 is an estimate of the average allele frequency (see above). We can then express the model in terms of the transformed sample covariance matrix:

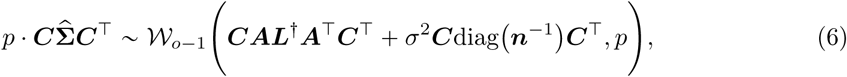

where 𝒲_*p*_ denotes a Wishart distribution with *p* degrees of freedom.^3^ Note we can equivalently use the sample squared Euclidean distance (often refereed to as a genetic distance) as a summary statistic: letting 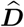 be the genetic distance matrix with 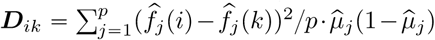, we have

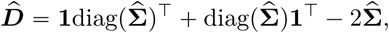

and so

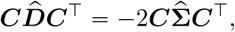

using the fact that the contrast matrix ***C*** is orthogonal to the one-vector. Thus we can use the same spatial covariance model implied by the allele frequencies once we project the distances on to the space of contrasts:^4^

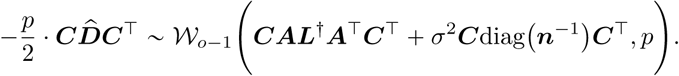

Overall, the negative log-likelihood function implied by our spatial model is (ignoring constant terms):

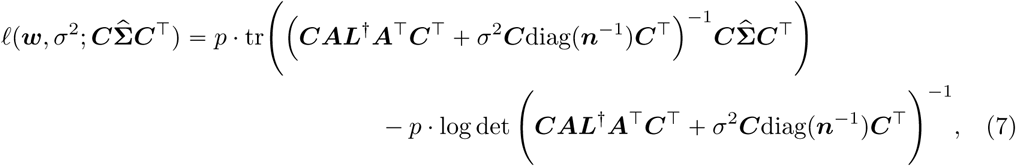

where ***w ∈****R*^*m*^ is a vectorized form of the non-zero lower-triangular entries of the weighted adjacency matrix ***W*** (recall that the graph Laplacian is completely defined by the edge weights ***L =*** diag (***W* 1) − *W*** so there is an implicit dependency here). Since the graph is a triangular lattice, we only need to consider the non-zero entries to save computational time, i.e. not all sub-populations are connected to each other.

One key difference between EEMS (Petkova et al., 2016) and FEEMS is how the edge weights are parameterized. In EEMS, each node is given an effective migration parameter *m*_*i*_ for node *i∈ 𝒱* and the edge weight is paramertized as the average between the nodes it connects, i.e. *w*_*ij*_ *=* (*m*_*i*_ *m*_*j*_)/2 for (*i, j*)∈ℰ. FEEMS, on the other hand, assigns a parameter to every nonzero edge-weight. The former has fewer parameters, with the specific consequence that it only allows isotropy and imposes an additional degree of similarity among edge weights; instead, in the latter, the edge weights are free to vary apart from the regularization imposed by the penalty. See Supp. Note “*Edge versus node parameterization*” and Supp. Fig. 16 for more details.

### Penalty description

As mentioned previously we would like to encourage that nearby edge weights on the graph have similar values to each other. This can be performed by penalizing the squared differences between all edges connected to the same node, i.e. spatially adjacent edges:

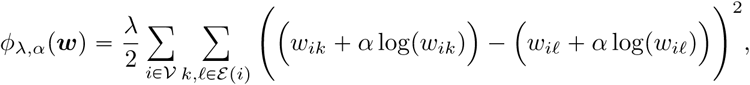

where *ϕ*_*λ,α*_ is our penalty function that represents the total amount of smoothness on the graph and ℰ(*i)* denotes the set of edges that connected to node *i*. Here we penalize a weighted combination of the edge weights on the original scale and logarithmic-scale where *α*, a tuning parameter, controls how strong the penalization is placed on the logarithmic scale—in the special case that *α =* 0, it reduces to the commonly used Laplacian smoothing-type penalty. Adding a logarithmic scale leads to smooth graphs for small edge values and thus allow for an additional degree of flexibility across orders of magnitude of edge weights. The smoothness parameter, *λ*, controls the overall contribution of the penalty to the objective function. It is convenient to write the penalty in matrix-vector form which we will use throughout:

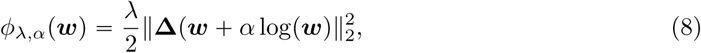

where **Δ** is a signed graph incidence matrix derived from a unweighted graph denoting if pairs of edges are connected to the same node. This penalty function (8) is also scale invariant, in the sense that for any 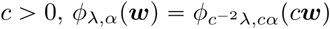.

One might wonder whether it is possible to use the *ℓ*_1_ norm in the penalty form (8) in place of the *ℓ*_2_ norm. While it is known that the *ℓ*_1_ norm might increase local adaptivity and better capture the sharp changes of the underlying structure of the latent allele frequencies, (e.g. Wang et al., 2016), in our case, we found an inferior performance when using the *ℓ*_1_ norm over the *ℓ*_2_ norm—in particular, our primary application of interest is the regime of highly missing nodes, i.e. *o ≪ d*, in which case the global smoothing seems somewhat necessary to encourage stable recovery of the edge weights at regions with sparsely observed nodes (see Supp. Note “*Smooth penalty with l*_1_ *norm*”). In addition, adding the penalty *ϕ*_*λ,α*_ (***w)*** allows us to implement faster algorithms to solve the optimization problem due to the differentiability of the *ℓ*_2_ norm, and as a result, it leads to better overall computational savings and a simpler implementation.

### Optimization

Putting (7) and (8) together, we infer the migration edge weights *ŵ* by minimizing the following penalized negative log-likelihood function:

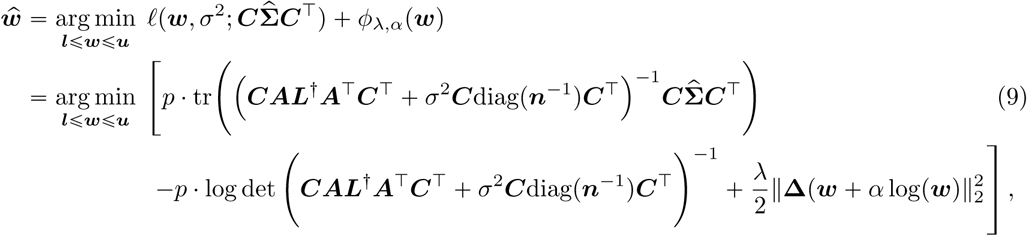

where 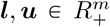 represent respectively the entrywise lower- and upper bounds on ***w***, i.e. we constrain the lower- and upper bound of the edge weights to ***l*** and ***u*** throughout the optimization. When no prior information is available on the range of the edge weights, we often set ***l*** = **0** and ***u*** = +∞.

One advantage of the formulation of (9) is the use of the vector form parameterization 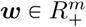 of the symmetric weighted adjacency matrix 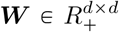. In our triangular graph *𝒢 =* (𝒱, ℰ), the number of non-zero lower-triangular entries is *m* = *𝒪 (d) ≪ d*^2^, so working directly on the space of vector parameterization saves computational cost. In addition, this avoids the symmetry constraint imposed on the adjacency matrix ***W***, hence making optimization easier (Kalofolias, 2016).

We solve the optimization problem using a constrained quasi-Newton optimization algorithm, specifically L-BFGS implemented in scipy (Byrd et al., 1995, Virtanen et al., 2020).^5^ Since our objective (9) is non-convex, the L-BFGS algorithm is guaranteed to converge only to a local minimum. Even so, we empirically observe that local minima starting from different initial points are qualitatively similar to each other across many datasets. The L-BFGS algorithm requires gradient and objective values as inputs. Note the naive computation of the objective (9) is computationally prohibitive since inverting the graph Laplacian has complexity 𝒪(*d*^3^). We take advantage of the sparsity of the graph and specific structure of the problem to efficiently compute gradient and objective values. In theory, our implementation has computational complexity of 𝒪(*do* + *o*^3^) per iteration which, in the setting of *o* ≪ *d*, is substantially smaller than 𝒪(*d*^3^).^6^

### Estimating the residual variance and edge weights under the null model

For estimating the residual variance parameter *σ*^2^, we first estimate it via maximum likelihood assuming homogeneous isolation by distance. This corresponds to the scenario where every edge-weight in the graph is given the exact same unknown parameter value *w*_0_. Under this model we only have two unknown parameters *w*_0_ and the residual variance *σ*^2^. We estimate these two parameters by jointly optimizing the marginal likelihood using a Nelder-Mead algorithm implemented in scipy (Virtanen et al., 2020). This requires only likelihood computations which are efficient due to the sparse nature of the graph. This optimization routine outputs an estimate of the residual variance 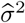 and the null edge weight ŵ_0_, which can be used to construct ***W***(*ŵ*) and in turn ***L***(*ŵ*_0_).

One strategy we found effective is to fit the model of homogeneous isolation by distance and then fix the estimated residual variance 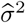 throughout later fits of the more flexible penalized models—See Supp. Note “*Jointly estimating the residual variance and edge weights*”. Additionally we find that initializing the edge weights to *ŵ*_0_ to be a useful and intuitive strategy to set the initial values for the entries of ***w*** to the correct scale.

### Data description and quality control

We analyzed a population genetic dataset of North American gray wolves previously published in Schweizer et al. (2016). For this, we downloaded plink formatted files and spatial coordinates from https://doi.org/10.5061/dryad.c9b25. We removed all SNPs with minor allele frequency less than 5% and with missingness greater then 10% resulting in a final set of 111 individuals and 17, 729 SNPs.

### Population structure analyses

We fit the Pritchard, Donnelly, and Stephens model (PSD) and ran principal components analysis on the genotype matrix of North American gray wolves (Price et al., 2006, Pritchard et al., 2000). For the PSD model we used the ADMIXTURE software on the un-normalized genotypes, running 5 replicates per choice of *K*, from *K =* 2 to *K =* 8 (Alexander et al., 2009). For each *K* we choose the one that achieved the highest likelihood to visualize. For PCA, we centered and scaled the genotype matrix and then ran sklearn implementation of PCA, truncated to compute 50 eigenvectors.

### Grid construction

To create a dense triangular lattice around the sample locations, we first define an outer boundary polygon. As a default, we construct the lattice by creating a convex hull around the sample points and manually trimming the polygon to adhere to the geography of the study organism and balancing the sample point range with the extent of local geography using the following website https://www.keene.edu/campus/maps/tool/. We often do not exclude internal “holes” in the habitat (e.g. water features for terrestrial animals), and let the model instead fit effective migration rates for those features to the extent they lead to elevated differentiation. We also emphasize the importance of defining the lattice for FEEMS as well as EEMS and suggest this should be carefully curated with prior biological knowledge about the system.

To ensure edges cover an equal area over the entire region we downloaded and intersected a uniform grid defined on the spherical shape of earth (Sahr et al., 2003). These defined grids are pre-computed at a number of different resolutions, allowing a user to test FEEMS at different grid densities which is an important feature to explore.

## Code Availability

The code to reproduce the results of this paper and more can be found in https://github.com/jhmarcus/feems-paper. A python package implementing the method can be found in https://github.com/jhmarcus/feems with documentation found in http://jhmarcus.com/feems/.

## Data Availability

We included a processed version of the dataset used in this manuscript in the feems package found here: https://github.com/jhmarcus/feems. An example tutorial on how to access the data the can be found here: http://jhmarcus.com/feems/notebooks/getting-started.html.

## Acknowledgements

We thank Rena Schweizer for helping us download and process the gray wolf dataset used in the paper, Ben Peter for providing feedback and code for helping to construct the discrete global grids and preparing the human genetic dataset, and Hussein Al-Asadi, Peter Carbonetto, Dan Rice for helpful conversations about the optimization and modeling approach. We also acknowledge helpful feedback from Arjun Biddanda, Anna Di Rienzo, Matthew Stephens, the Stephens Lab, the Novembre Lab, and the University of Chicago 4th floor computational biology group.

This study was supported in part by the National Science Foundation via fellowship DGE-1746045 and the National Institute of General Medical Sciences via training grant T32GM007197 to J.H.M and R01GM132383 to J.N. W.H. was partially supported by the NSF via the TRIPODS program and by the Berkeley Institute for Data Science. R.F.B. was supported by the National Science Foundation via grant DMS–1654076, and by the Office of Naval Research via grant N00014-20-1-2337.

## Author Contributions

J.H.M and J.N. conceived of the project. J.H.M. and W.H. developed the statistical methodology with guidance from R.F.B. (lead) and J.N. (supporting). J.H.M, W.H. carried out method testing and application with guidance from J.N. (lead) and R.F.B. (supporting). J.H.M. and W.H. developed the software. J.H.M. and W.H. wrote the paper with edits from R.F.B. and J.N.

## Supplementary Materials

### Mathematical notation

We denote matrices using bold capital letters ***A***. Bold lowercase letters are vectors ***a***, and non-bold lowercase letters are scalars *a*. We denote by ***A***^−1^ and ***A^†^*** the inverse and (Moore-Penrose) pseudo-inverse of ***A*** respectively. We use ***y ∼****N*_*p*_ (***µ*, Σ)** to express that the random vector ***y*** is modeled as a *p*-dimensional multivariate Gaussian distribution with fixed parameters ***µ*** and **Σ** and use the conditional notation ***y***|***µ ∼*** *N*_*p*_(***µ*, Σ**) if ***µ*** is random.

A graph is a pair *𝒢 =* (𝒱, ℰ), where 𝒱 denotes a set of nodes or vertices and *ℰ* ⊂ 𝒱 × 𝒱 denotes a set of edges. Throughout we assume the graph *𝒢* is undirected, weighted, and contains no self loops, i.e. (*i, j*) ∈ *ℰ⇔*(*j, i*) *∈ ℰ* and = (*i, i*) ∉ *ℰ* and each edge (*i, j*) ∈ *ℰ* is given a weight *w*_*ij*_ = *w*_*ji*_ > 0. We write ***W*** to indicate the symmetric weighted adjacency matrix, i.e.

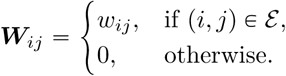

***w ∈*** *R*^*m*^ is a vectorized form of the non-zero lower-triangular entries of ***W*** where *m =* |*ℰ*|*/*2 is the number of non-zero lower triangular elements. We denote by ***L =*** diag (***W* 1) − *W*** the graph Laplacian.

### Gradient computation

In practice, we make a change of variable from 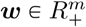 to *z* = log(*w*) ∈ *R^m^* and the algorithm is applied to the transformed objective function:

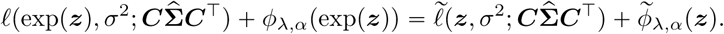

After the change of variable, the objective value remains the same whereas it follows from the chain rule that 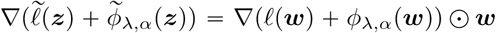 where ⊙ indicates the Hadamard product or elementwise product—for notational convenience, we drop the dependency of *ℓ* on the quantities *σ*^2^ and 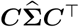. Furthermore, the computation of ∇*ϕ*_*λ,α*_ (***w)*** is relatively straight-forward, so in the rest of this section, we discuss only the computation of the gradient of the negative log-likelihood function with respect to ***w***, i.e. ∇*ℓ(****w)***.

Recall, by definition, the graph Laplacian ***ℓ*** implicitly depends on the variable ***w*** through ***L =*** diag (***W* 1) − *W***. Throughout we assume the first *o* rows and columns of ***L*** correspond to the observed nodes. With this assumption, our node assignment matrix has block structure ***A = [*I**_*o*×*o*_ |**0**_*o*×(*d*−*o*)_]. To simplify some of the equations appearing later, we introduce the notation: we define

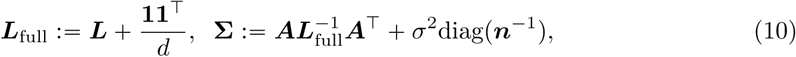

and

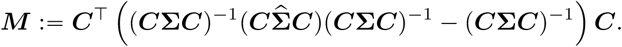

Applying the chain rule and matrix derivatives, we can calculate:

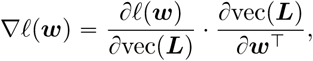

where vec is the vectorization operator and ∂*ℓ/*∂vec(***L***) and ∂vec(***L***)/ ∂***w***^T^ are 1 × *d*^2^ vector and *d*^2^ × *d* matrix, respectively, given by

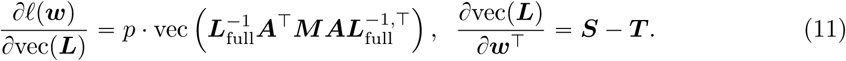

Here ***S*** and ***T*** are linear operators that satisfy ***Sw =*** diag (***W* 1)** and ***T w = W***. Note ***S*** and ***T*** both have 𝒪(*d*) many nonzero entries, so we can perform sparse matrix multiplication to efficiently compute the matrix-vector multiplication ∂*l/*∂ vec (***L)·(S−T)***. On the other hand, the computation of ∂*l/*∂ vec (***L)*** is more challenging as it requires inverting the full *d ×d* matrix ***L***_full_. Next we develop a procedure that efficiently computes ∂*l/* ∂vec (***L)***. We proceed by dividing the task into multiple steps.

1. **Computing Σ**^−1^ Recalling the block structure ***A* [I**_*o*×*o*_ |**0**_*o*×(*d*−*o*)_] of the node assignment matrix, we can write **Σ** as:

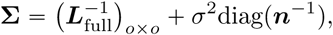

where 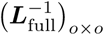 denotes the *o* × *o* upper-left block of 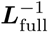. Following Petkova et al. (2016), the inverse **Σ**^−1^ has the form

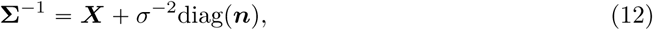

for some matrix ***X*** ∈ *R*^*o*×*o*^. Equating **ΣΣ**^−1^ = **I**, it follows that

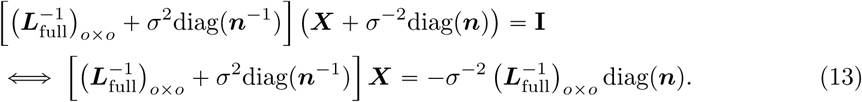 Therefore, **Σ**^−1^ can be obtained by solving the *o*×*o* linear system (13) and plugging the solution into (12). The challenge here is to compute 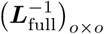 without matrix inversion of the full-dimensional ***L***_***full***_
2. **Computing** 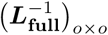 Let ***L***_full,*o*×*o*_ be the *o* × *o* block matrix corresponding to the observed nodes of ***L***_full_ and and similarly let ***L***_full,(*d*−*o*)×(*d*−*o*)_ and 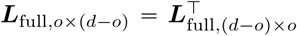 be the corresponding block matrices of ***L***_full_ respectively. The inverse of 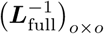 is then given by the Schur complement of ***L*** _full,(*d*−*o*)×(*d*−*o*)_ in ***L***:

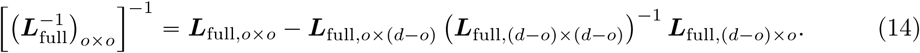 See also Hanks and Hooten (2013), Petkova et al. (2016). Since every term in (14) has sparse + rank-1 structure, the matrix multiplications can be performed fast. In addition, for the term (***L*** _full,(*d*−*o*)×(*d*−*o*)_) ^−1^,we can use the Sherman-Morrison formula so that the inverse is given explicitly by

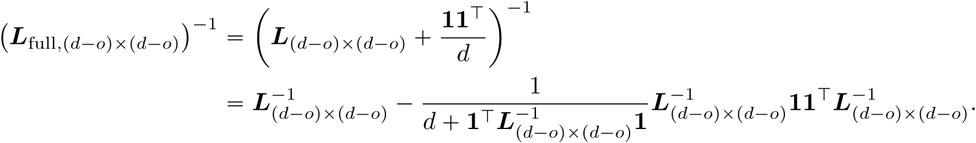 Hence, in order to compute (***L*** _full,(*d*−*o*)×(*d*−*o*)_) ^−1^ ***L*** _full,(*d*−*o*)×*o*_, we need to solve two systems of linear equations:

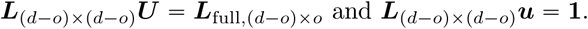 Note that the matrix ***L***_(*d*−*o*)×(*d*−*o*)_ is sparse, so both systems can be solved efficiently by performing sparse Cholesky factorization on ***L***_(*d*−*o*)×(*d*−*o*)_ (Hanks and Hooten, 2013). Alternatively, one can implement fast Laplacian solvers (Vishnoi et al., 2013) that solve the Laplacian system in time nearly linear in the dimension *𝒪*(*d*). After we obtain 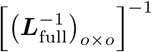 via sparse + rank-1 matrix multiplication and sparse Cholesky factorization, we can invert the *o* × *o* matrix to get 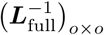.
3. **Computing** 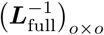 Write

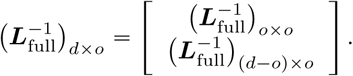 Using the inversion of the matrix in a block form, the (*d* −*o*) ×*o* block component is given by

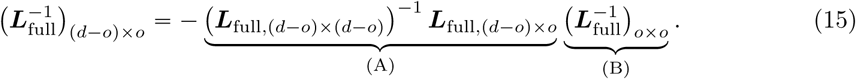 Since each of the two terms (A) and (B) has been already computed in the previous step, there is no need to recompute them. In total, it requires a (*d −o) × o* matrix and *o ×o* matrix multiplication.
4. **Computing the full gradient** Going back to the expression of *∇ℓ (****w)*** in (11), and noting the block structure of the assignment matrix ***A***, we have:

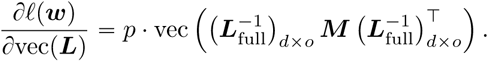 Let Π_**1**_ **1 (1**^**T**^**Σ**^**−**T^**1)** ^**−**T^ **1**^T^**Σ**^**−**T^ be projection to the space of constant vectors with respect to the inner product ⟨***x, y***⟩= ***x***^T^**Σ**^**−**T^ ***y***. Using the identity **I Π**_**1**_ **= Σ*C***^T^ (***C*Σ*C***^T^) ^**−**T^ ***C*** (McCullagh, 2009), then we can write ***M*** in terms of **Π**_**1**_:

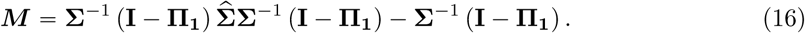 Since Π_**1**_ is a rank-1 matrix, this expression of ***M*** allows easier computation. Finally we can put together (12), (13), (15), and (16), to compute the gradient of the negative log-likelihood function with respect to the graph Laplacian.

### Objective computation

The graph Laplacian ***L*** is orthogonal to the one vector **1**, so using the notation introduced in (10), we can express our objective function as

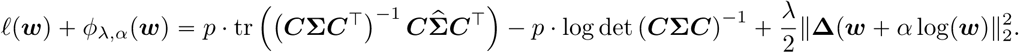

With the identity **I** − **Π**_**1**_ = **Σ*C***^T^(***C*Σ*C***^T^) ^**−**T^ ***C***, the trace term is:

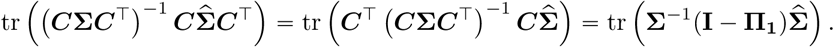

The matrix inside the trace has been constructed in the gradient computation, see equation (16). In terms of the determinant, we use the same approach considered in Petkova et al. (2016)—in particular, concatenating ***C***^T^ and **1**, the matrix [***C***^T^ | 1] is orthogonal, so it can be shown that

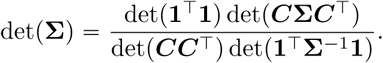

Rearranging terms and using the fact det(***U*** ^**−**T^) = det(***U***) ^**−**T^ for any matrix ***U***, we obtain:

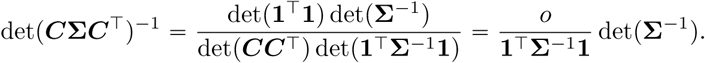

We have computed **Σ**^**−**T^ in equation (12), so each of the terms above can be computed without any additional matrix multiplications. Finally, the signed graph incidence matrix **Δ** defined on the edges of the graph is, by construction, highly sparse with 𝒪(*d)* many nonzero entries. Hence we implement sparse matrix multiplication to evaluate the penalty function *ϕ*_*λ,α*_ (***w)*** while avoiding the full-dimensional matrix-vector product.

### Estimating the edge weights under the exact likelihood model

Recall that, when describing our data model, we employed the approximation 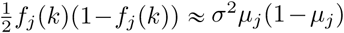 for all SNPs *j* and nodes *k* (see equation (4)) and estimated the residual variance *σ*^2^ under the homogeneous isolation by distance model. Here we examine whether this approximation results in a significant difference with respect to the estimation quality of the edge weights of the graph.

Without approximation, we can calculate the exact analytical form for the marginal likelihood of the estimated frequency as follows (after removing the SNP means):

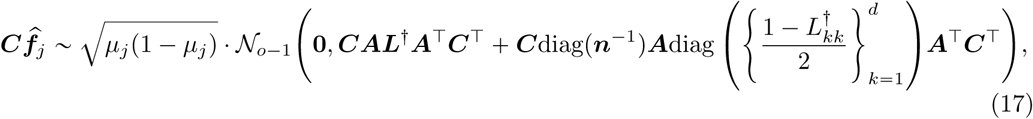

where 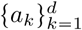 represents the vector ***a*** = (*a*_1_, *…, a*_*d*_). We then consider estimating the edge weights with the likelihood based on (17) and without relying on approximating the residual variance. In particular, comparing to the model (5), this formulation does not introduce the unknown residual variance parameter *σ*_2_ but rather it is given implicitly by 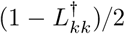. This means that the model (17) is well-defined only when 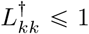 for all nodes *k*, hence leading to the following constrained optimization problem:

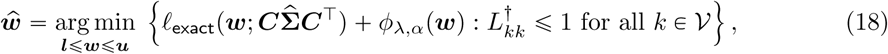

where ℓ_exact_ is the negative log-likelihood function implied by the model (17) and *ϕ*_*λ,α*_ is our smooth penalty function. The main difficulty of solving (18) is that enforcing the constraint 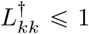 for all nodes *k*, ∈ 𝒱 requires full computation of the pseudo-inverse of a *d × d* matrix ***L*** whereas in order to evaluate the likelihood, we only need to calculate ***L^†^*** on the observed nodes. To overcome this computational challenge, we may relax the constraint and consider the following form as a proxy for optimization (18):

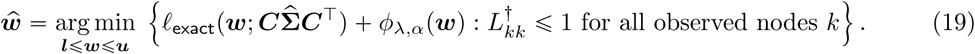

We can solve this problem efficiently using a gradient-based algorithm where the gradient of *l*_exact_ with respect to ***L*** is given by

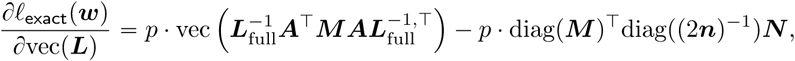

where ***M*** is a *o* × *o* matrix defined in (16), while ***N*** is a *o* × *d*^2^ matrix whose rows correspond to the observed subsets of the rows of the *d*^2^ × *d*^2^ matrix 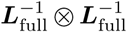.

Overall, when we implement the penalized restricted maximum likelihood procedure in (19), we find that it does not make much of a difference and output qualitatively comparable results to FEEMS—for example, Supp. Fig. 12 shows one such fit with a setting of *λ =* 10^−3^ and *α =* 50. Unfortunately, this approach has a drawback that after the algorithm reaches the solution, the term 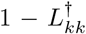 is not guaranteed to be positive for the unobserved nodes, since, due to the computational efficiency, the constraints 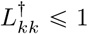 are only placed on the observed nodes. This, in principle, results in an ill-defined model if we would like interpretable results at unobserved as well as observed nodes, and therefore we replace the calculation (17) with the approximation (5) to avoid this issue. In addition, by decoupling the residual variance parameter *σ*^2^ from the graph-related weighted edges ***w***, the model (6) has more resemblance to spatial coalescent model used in EEMS (Petkova et al., 2016).

### Jointly estimating the residual variance and edge weights

One simple strategy we have used throughout the paper was to fit *σ*^2^ first under a model of homogeneous isolation by distance and prefix the estimated residual variance to the resulting 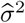 for later fits of the effective migration rates. Alternatively, one might come up with a strategy to estimate the unknown residual variance jointly with the edge weights, instead of prefixing it from the estimation of the null model—the hope here is to simultaneously correct the model misspecification and allow for improving model fit to the data.

As it turns out, given such a small fraction of sampled spatial locations in the data, the strategy of jointly optimizing the marginal likelihood with respect to both variables has the tendency to overfit to the data unless it is properly regularized. Specifically, we can consider the model that generalizes (6), namely

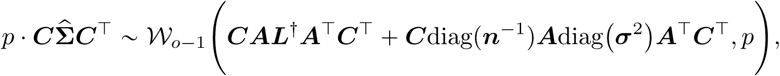

where ***σ***^2^ is a *d ×* 1 vector of node specific residual variances, i.e. each deme has its own residual parameter *σ*_*k*_ for all nodes *k*. If the node specific parameters *σ*_*k*_’s are assumed to be same across all nodes, this reduces to the model (6). Supp. Fig. 13 shows the results of different strategies of estimating the residual variances. As expected, when the model has a single residual variance *σ*^2^, either prefixing it from the null model (Figure 4) or estimating it jointly with the edge weights (Supp. Fig. 13A) lead to similar and comparable outputs. The major difference is the high migration edge forming long path appearing in Supp. Fig. 13A to separate the reduced gene-flows in the middle, which tends to disappear as *α* increases. Whereas, if the residual variances are allowed to be node specific, the fitted 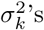 are highly variable and as a result the estimated graph misses some geographic features present in the data, such as reduced effective migration around St. Lawerence Island (Supp. Fig. 13B). Presumably this is attributed to overfitting, due to the absence of data in many unobserved demes. In EEMS, in order to estimate the genetic diversity parameters for every spatial position, which play a similar role as the residual variance in FEEMS, a Voronoi-tessellation prior is placed to encourage sharing of information across adjacent nodes and prevent over-fitting. While we can similarly estimate the node specific residual variances on every node of the graph with our penalty function (*ϕ*_*λ,α*_ defined on the variable ***σ***^2^, we do not find it substantially improves the extent to which the model suits the data. Thus, we take the approach of fitting the single residual variance *σ*^2^ under the null model and prefixing it as a simple but effective strategy with apparent good empirical performance.

### Edge versus node parameterization

One of the novel features of FEEMS is its ability to directly find the edge weights of the graph that best suit the data. This direct edge parameterization may increase the risk of model’s overfitting, but also allows for more flexible estimation of migration histories. Furthermore, as seen in Figure 2 and Supp. Fig. 2, it has potential to recover anisotropic migration processes. This is in contrast to EEMS wherein every spatial node is assigned an effective migration parameter *m*_*k*_ and a migration rate on each edge joining nodes *k* and *k*′ is given by the average effective migration *w*_*kk*′_ = (*m*_*k*_ *m*_*k*′_)/2. Not surprisingly, parameterization via node-specific parameters induces implicit regularization by substantially constraining the feasible set of graph’s edge weights. In some cases, this has the desirable property of imposing an additional degree of similarity among edge weights, but it often restricts the model’s capacity to capture a richer set of structure present in the data, (e.g. Petkova et al., 2016, supplementary figure 2). To be concrete, Supp. Fig. 15 displays two different fits of FEEMS based on edge parameterization (Supp. Fig. 15A) and node parameterization (Supp. Fig. 15B), run on a previously published dataset of human genetic variation from Africa (see Peter et al. (2018) for details on the description of the dataset). Running FEEMS with a node-based parameterization is straightforward in our framework—all we have to do is to reparameterize the edge weights by the average effective migration and solve the corresponding optimization problem (9) with respect to ***m***. It is evident from the results that FEEMS with edge parameterization exhibits subtle correlations that exist between the annotated demes in the figure whereas node parameterization fails to recover them. We also compare the model fit of FEEMS to the observed genetic covariance (Supp. Fig. 16) and find that edge-based parameterization provides a better fit to the African dataset. Supp. Fig. 17 further demonstrates that in the coalescent simulations with anisotropic migration, the node parameterization is unable to recover the ground truth of the underlying migration rates even when the nodes are fully observed.

### Smooth penalty with ℓ_1_ norm

FEEMS’s primary optimization objective (see equation (9)) is:

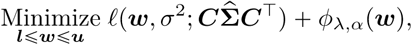

where the spatial smoothness penalty is given by 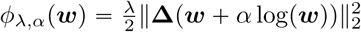. It is widely known that ℓ_1_-based method leads to better local adaptive fitting and structural recovery than ℓ_2_-based methods (Wang et al., 2016), but at the cost of handling non-smooth objective functions that are often computationally more challenging and demanding. In a spatial genetic dataset, one major challenge is to deal with the relatively sparse sampling design where there are many unobserved nodes on the graph. In this challenging statistical setting, our finding is that an ℓ_2_-based method enables more accurate and reliable estimation of the geographic features.

Specifically, writing 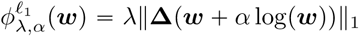, we considered the alternate following composite objective function:

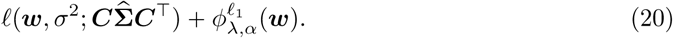

To solve (20), we apply linearized alternating direction method of multipliers (ADMM) (Boyd et al., 2011), a variant of the standard ADMM algorithm, that iteratively optimizes the augmented Lagrangian over the primal and dual variables. The derivation of the algorithm is a standard calculation so we omit the detailed description of the algorithm. As opposed to the common belief about the effectiveness of the ℓ_1_ norm for structural recovery, the recovered graph of FEEMS using ℓ_1_-based smooth penalty shows less accurate reconstruction of the migration patterns, particularly when the sampling design has many locations with missing data on the graph (Supp. Fig. 18A, Supp. Fig. 19H). It appears that the ℓ_1_-based penalty function is not capable of accurately estimating edge weights at regions with little data, partially due to its local adaptation, in contrast to the ℓ_2_-based method that considers regularization more globally. This suggests that in order to use the ℓ_1_ penalty 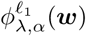 in the presence of many missing nodes, one needs an additional regularization term that promotes global smoothness of the graph’s edge weights, e.g., a combination of 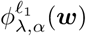 and *ϕ*_*λ,α*_(***w***) (same spirit as elastic net (Zou and Hastie, 2005)), or 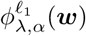 on top of node-based parameterization (Supp. Fig. 18B).

## Supplementary Figures

**Supplementary Figure 1:**
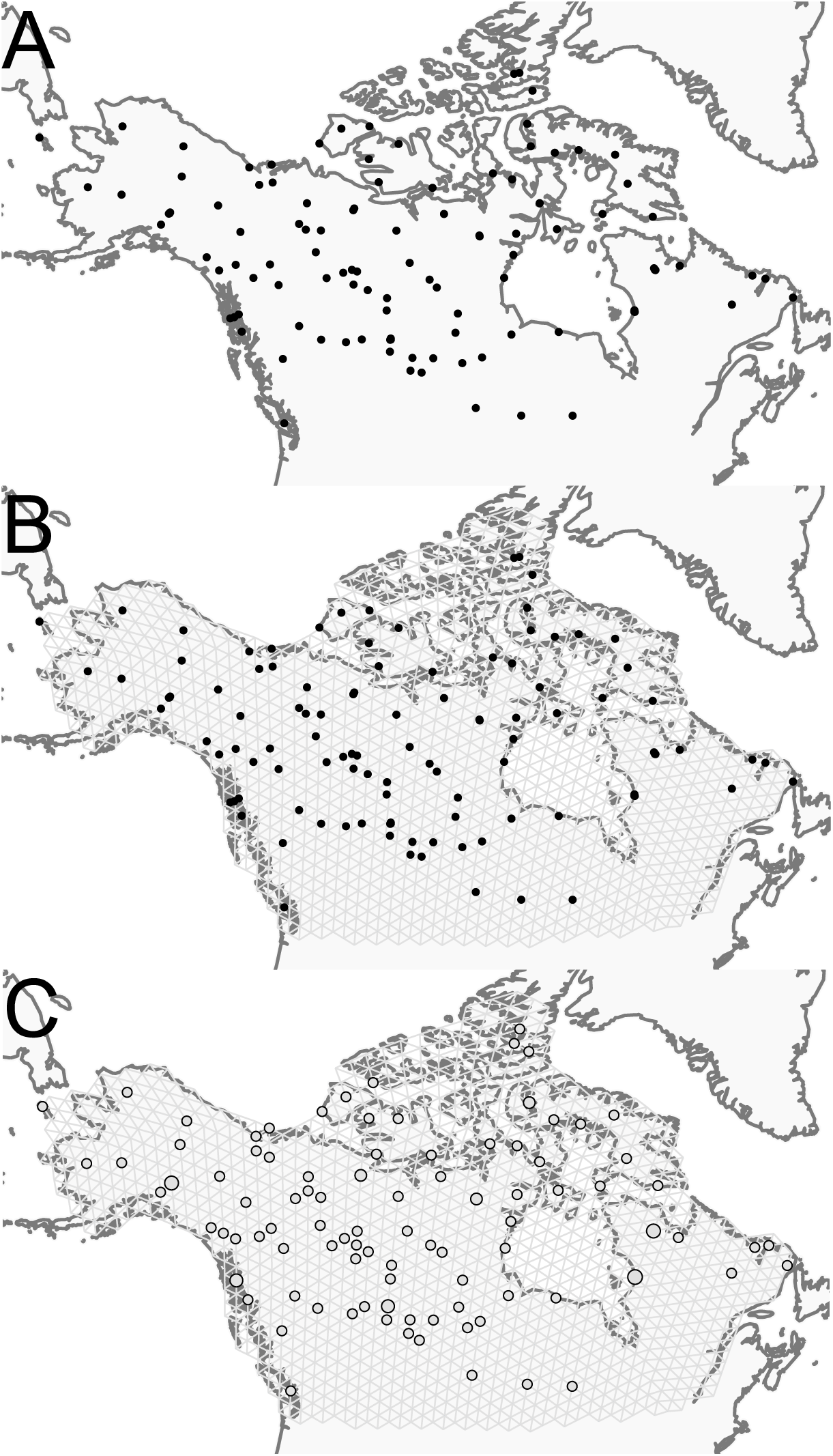
Visualization of grid construction and node assignment: (A) Map of sample coordinates (black points) from a dataset of gray wolves from North America. The input to FEEMS are latitude and longitude coordinates as well as genotype data for each sample. (B) Map of sample coordinates with an example dense spatial grid. The nodes of the grid represent sub-populations and the edges represent local gene-flow between adjacent sub-populations. (C) Individuals are assigned to nearby nodes (sub-populations) and summary statistics (e.g., allele frequencies) are computed for each observed location.

**Supplementary Figure 2:**
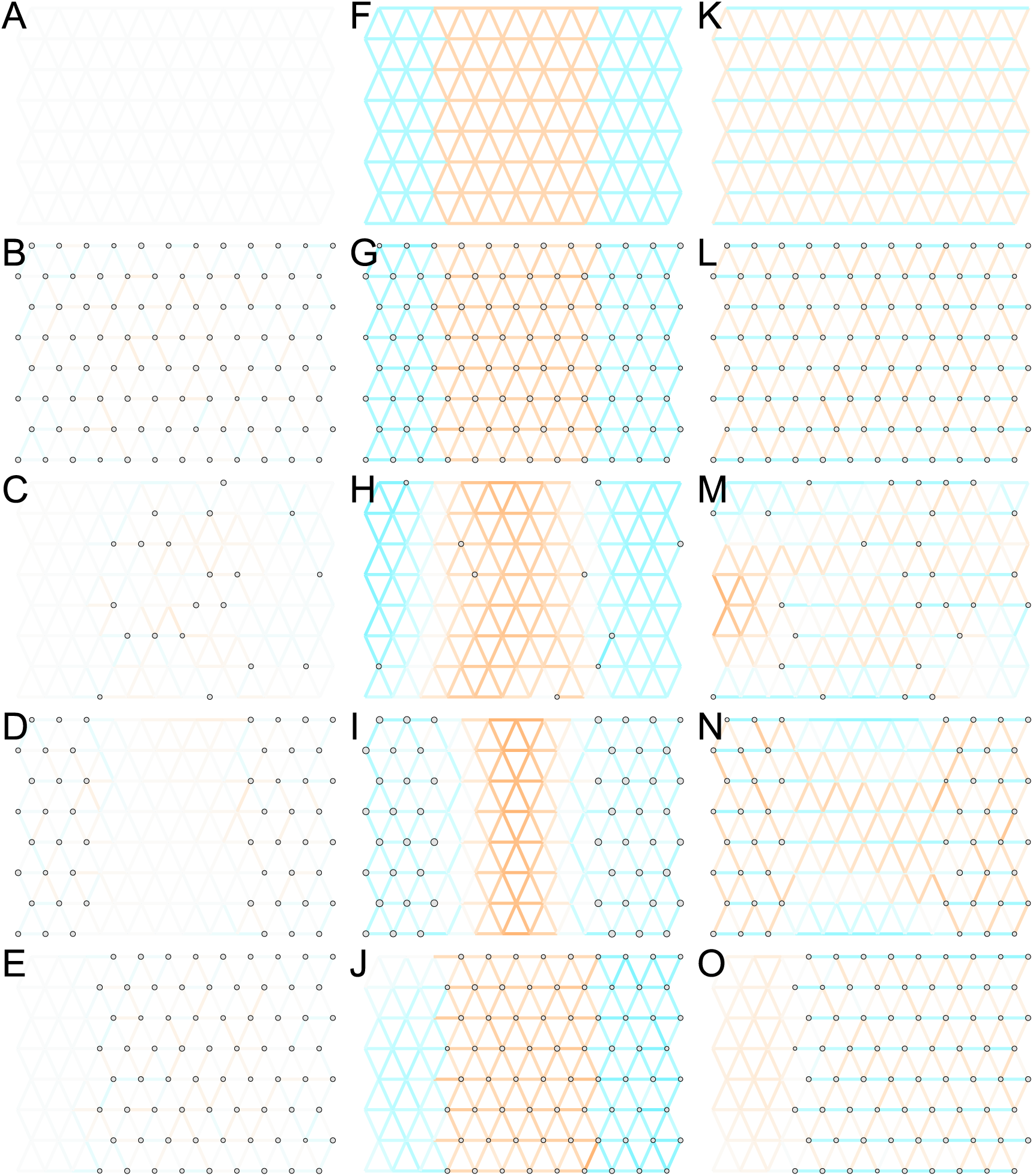
Application of FEEMS to an extended set of coalescent simulations: We display an extended set of coalescent simulations with multiple migration scenarios and sampling designs. The sample sizes across the grid are represented by the size of the grey dots at each node. The migration rates are obtained by solving FEEMS objective function (9) where the regularization parameters are specified at *λ =* 10^−2^, *α =* 30 (I), *λ* = 10^−4^, *α =* 30 (N), and *λ =* 10^−3^, *α =* 30 for the rest. (A, F, K) display the ground truth of the underlying migration rates. (B, G, L) Shows simulations where there is no missing data on the graph. (C, H, M) Shows simulations with sparse observations and nodes missing at random. (D, I, N) Shows simulations of biased sampling where there are no samples from the center of the simulated habitat. (E, J, O) Shows simulations of biased sampling where there are only samples on the right side of the habitat.

**Supplementary Figure 3:**
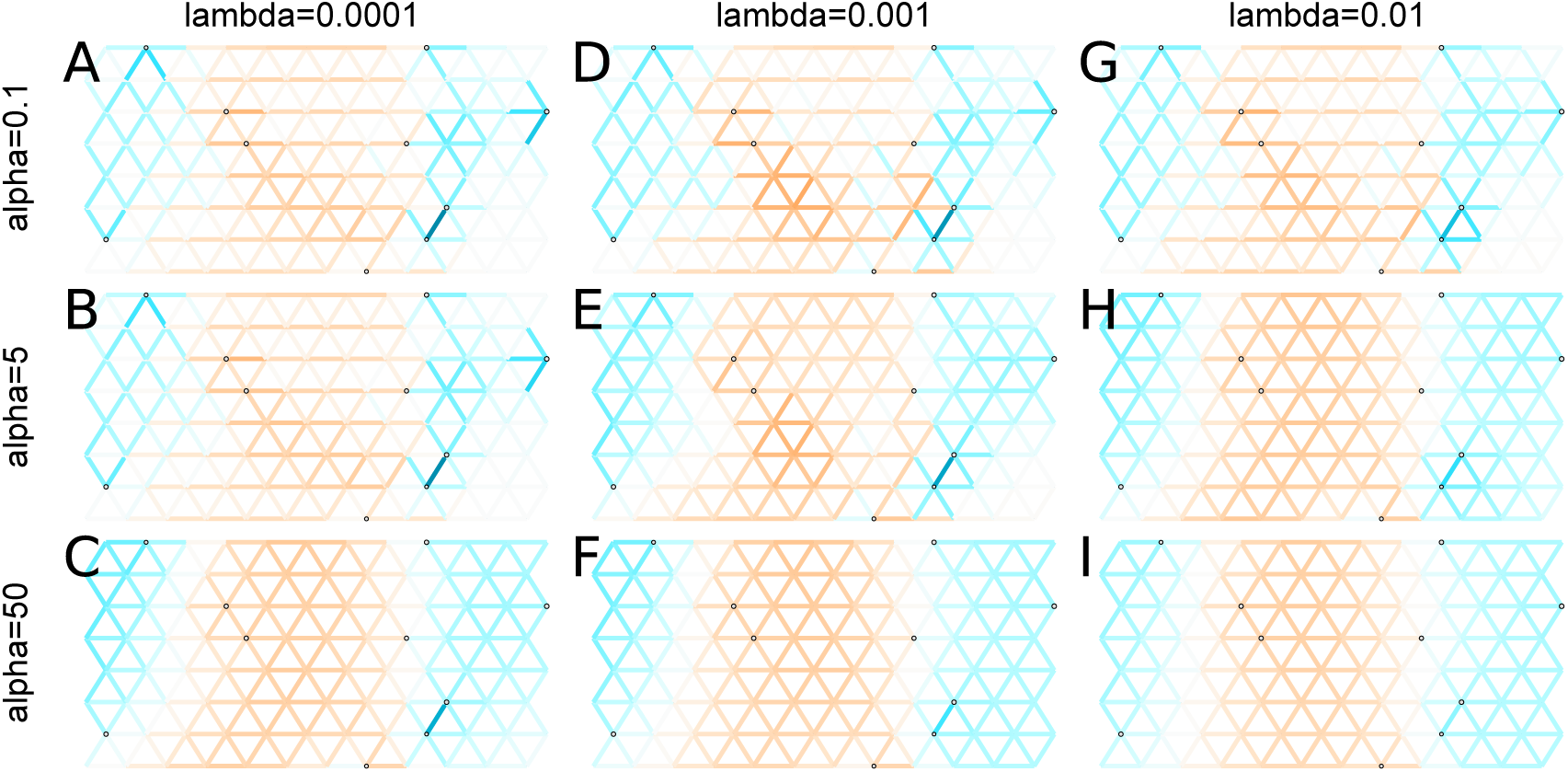
Application of FEEMS to a heterogeneous migration scenario with a “missing at random” sampling design: We run FEEMS on coalescent simulation with a non-homogeneous process while varying hyperparameters *λ* (rows) and *α* (columns). We randomly sample individuals for 20% of nodes. When *λ* grows, the fitted graph becomes overall smoother, whereas *α* effectively controls the degree of similarity among low migration rates.

**Supplementary Figure 4:**
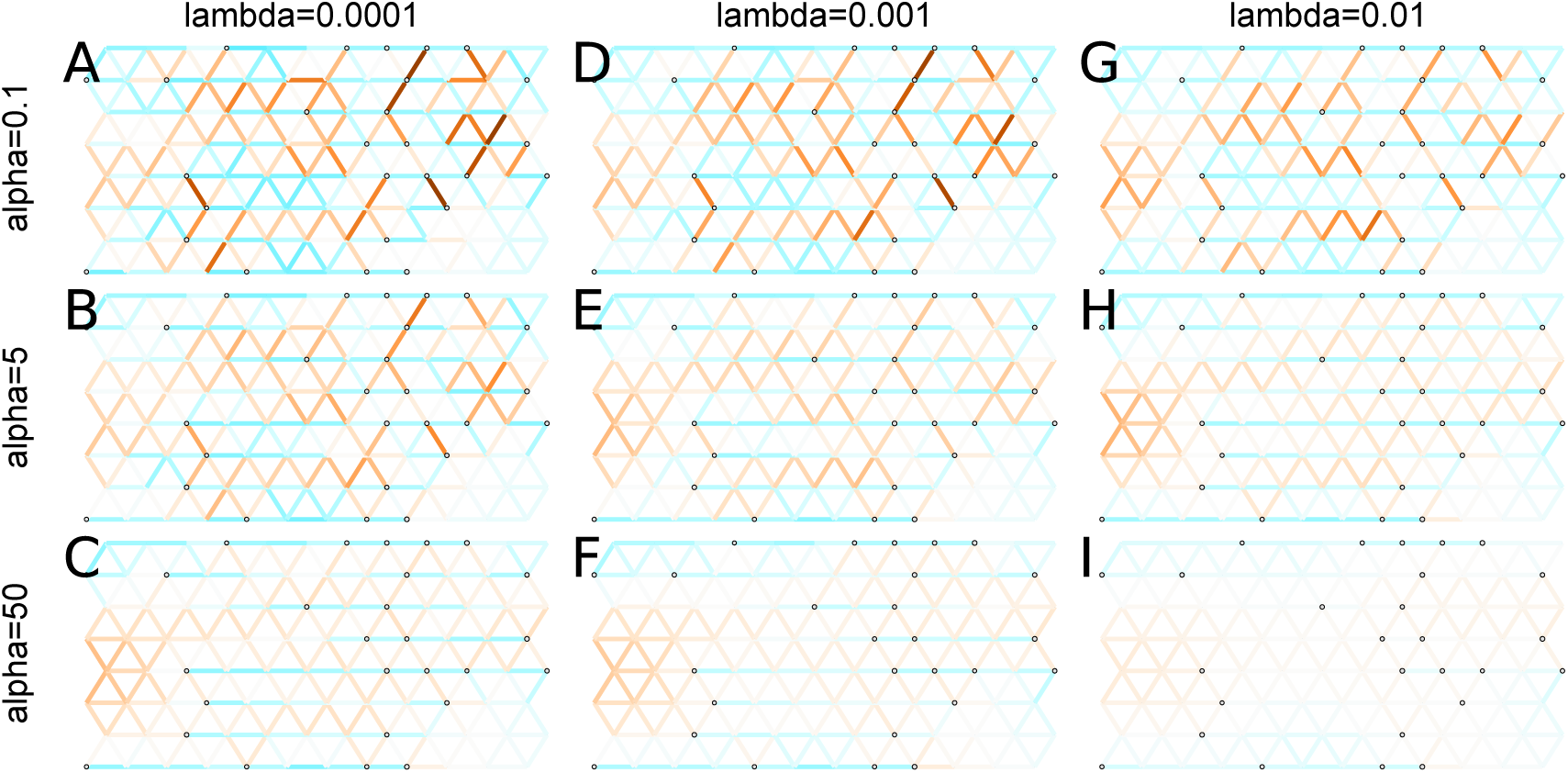
Application of FEEMS to an anisotropic migration scenario with a “missing at random” sampling design: We run FEEMS on coalescent simulation with an anisotropic process while varying hyperparameters *λ* (rows) and *α* (columns). We randomly sample individuals for 20% of nodes. When *λ* grows, the fitted graph becomes overall smoother, whereas *α* effectively controls the degree of similarity among low migration rates.

**Supplementary Figure 5:**
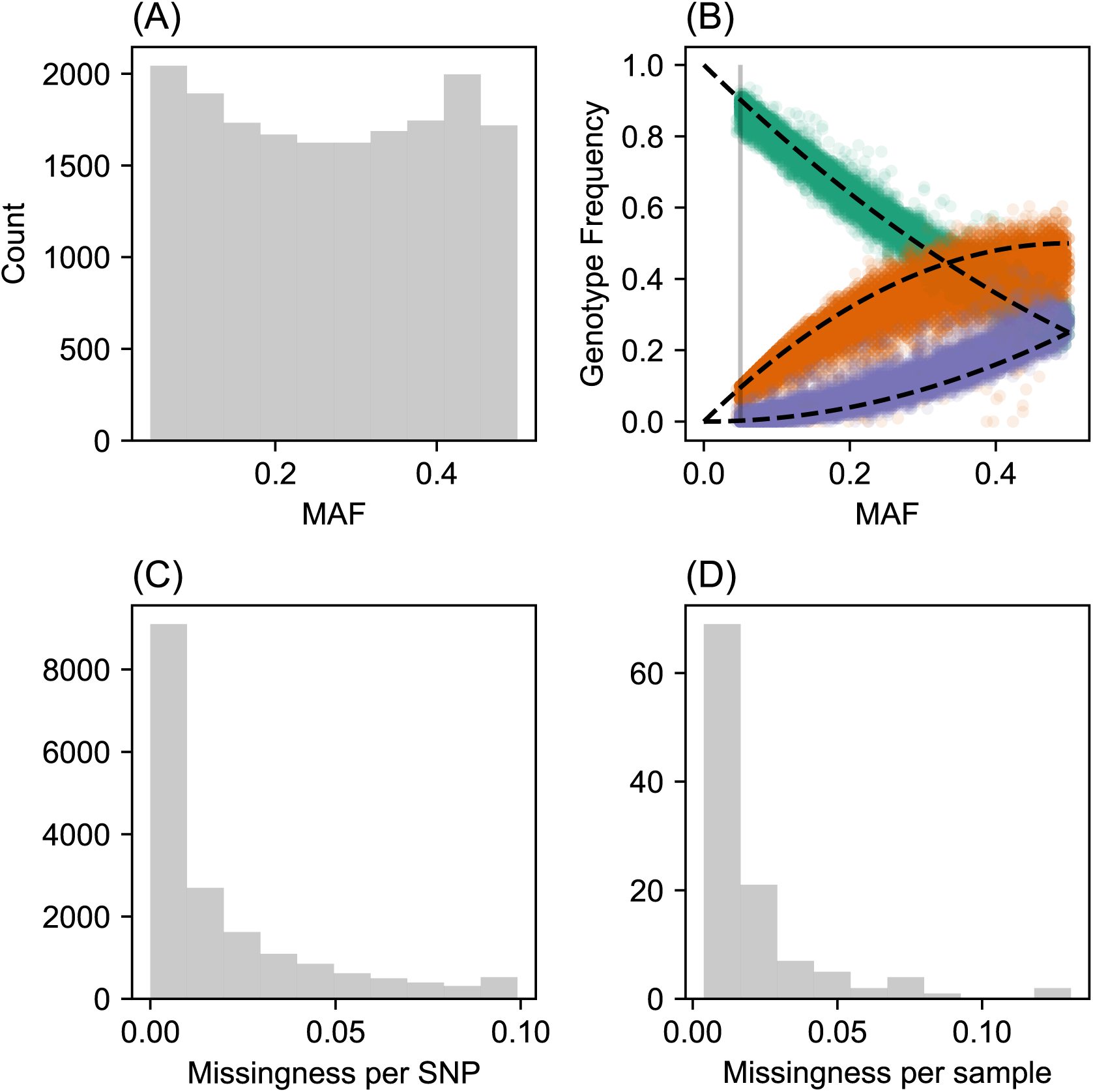
SNP and individual quality control: (A) Displays a visualization of the sample site frequency spectrum. Specifically, we display a histogram of minor allele frequencies across all SNPs. We see a relatively uniform histogram which reflects the ascertainment of common SNPs on the array that was designed to genotype gray wolf samples. (B) Visualization of allele frequencies plotted against genotype frequencies. Each point represents a different SNP and the colors represent the 3 possible genotype values. The black dashed lines display the expectation as predicted from a simple binomial sampling model i.e. Hardy-Weinberg equilibrium. (C) Displays a histogram of the missingness fraction per SNP. We observe the missingness tends to be relatively low for each SNP. (D) Displays a histogram of the missingness fraction per sample. Generally, the missingness tends to be low for each sample.

**Supplementary Figure 6:**
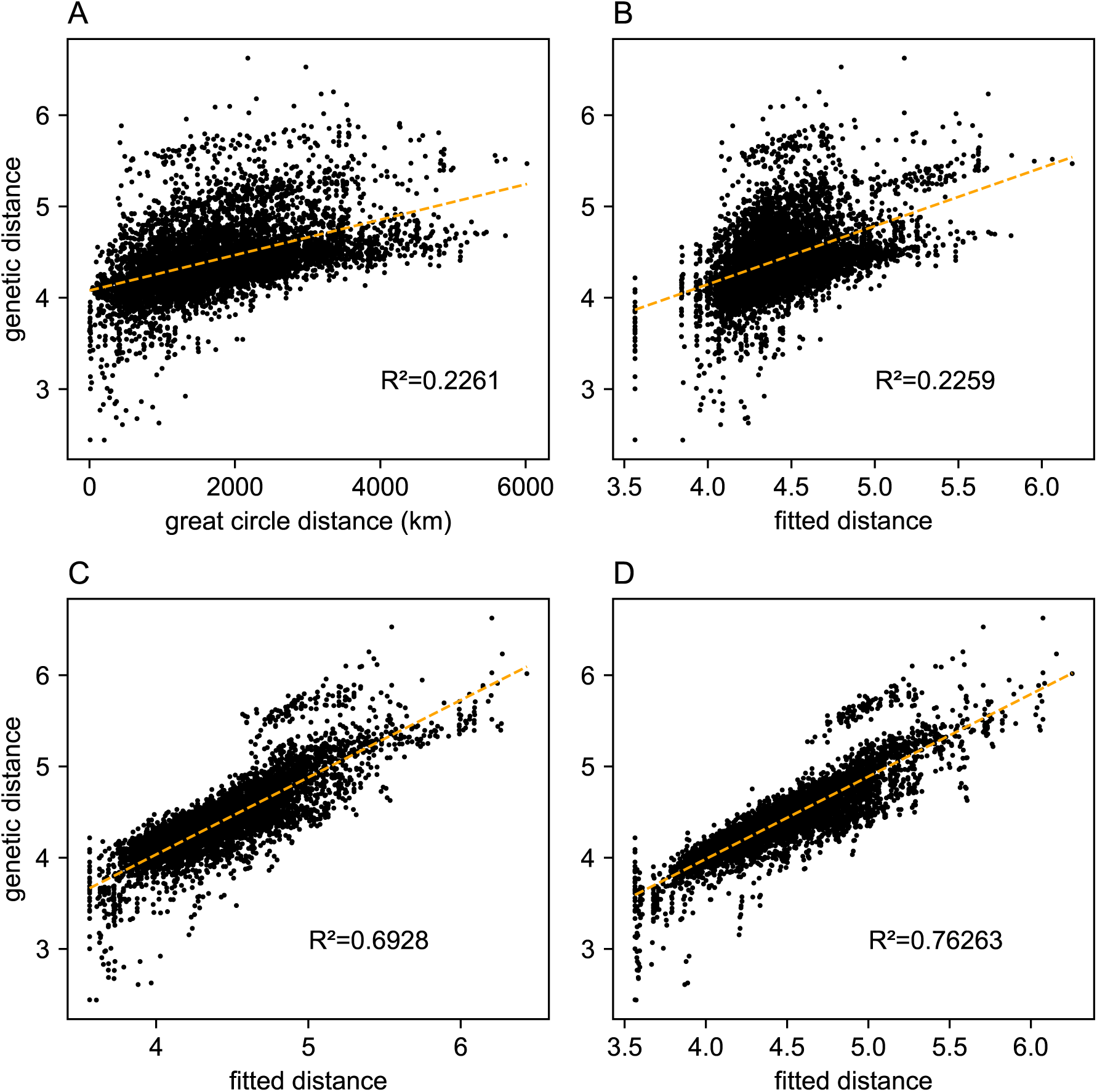
Comparing predictions of observed genetic distances: We display different predictions of observed genetic distances using geographic distance or the fitted genetic distance output by FEEMS. (A) The x-axis displays the geographic distance between two individuals, as measured by the great circle distance (haversine distance). The y-axis displays the squared Euclidean distance between two individuals averaged over all SNPs. (B-D) The x-axis displays the fitted genetic distance as predicted by the FEEMS model and y-axis displays the squared Euclidean distance between two individuals averaged over all SNPs. For (B-D) we display the fit of *λ* getting subsequently smaller (10, 10^−3^, 10^−5^)and as expected the fit becomes better because we tolerate more complex surfaces and we are not evaluating the fit on out-of-sample data.

**Supplementary Figure 7:**
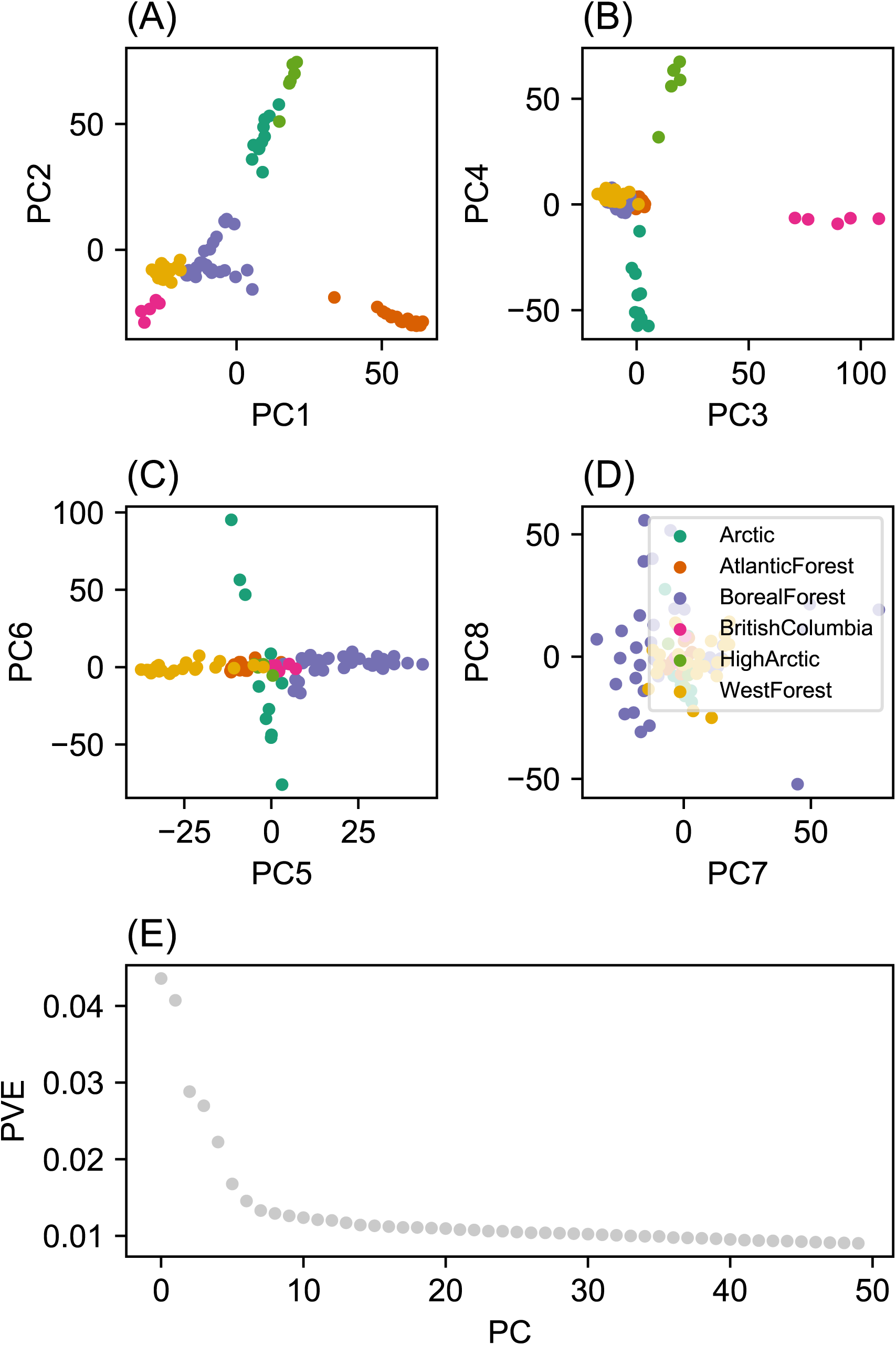
Summary of top axes of genotypic variation: We display a visual summary of Principal Components Analysis (PCA) applied to the normalized genotype matrix from the North American gray wolf dataset. (A-D) Displays PC bi-plots of the top seven PCs plotted against each other. The colors represent predefined ecotypes defined in (Schweizer et al., 2016). We can see that the top PCs delineate these predefined ecotypes. (E) Shows a “scree” plot with the proportion of variance explained for each of the top 50 PCs. As expected by genetic data (Patterson et al., 2006), the eigen-values of the genotype matrix tend to be spread over many PCs.

**Supplementary Figure 8:**
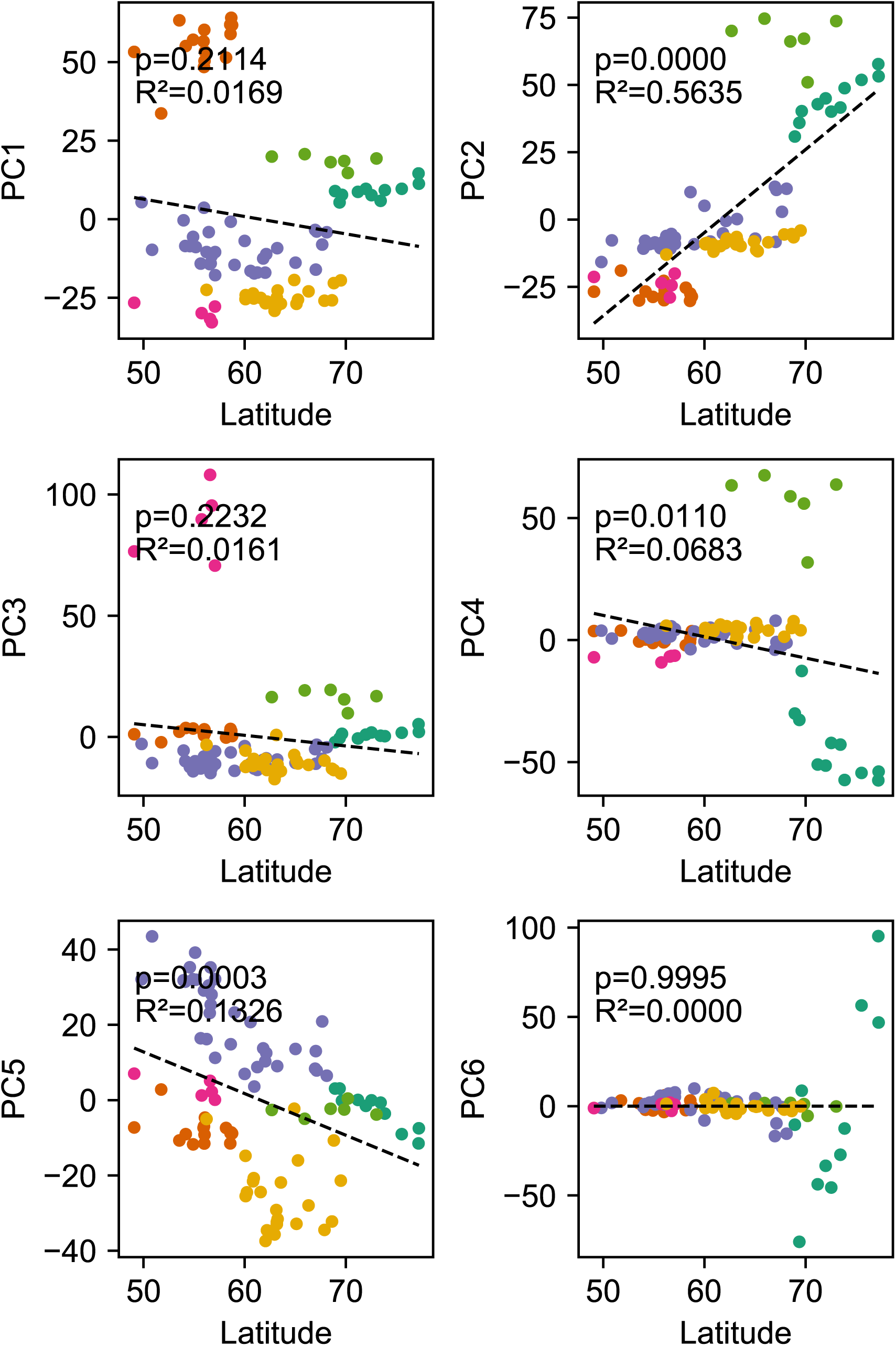
Relationship between top axes of genetic variation and latitude: In each sub-panel we plot the PC value against latitude for each sample in gray the wolf dataset. We see many of the top PCs are significantly correlated with latitude as tested by linear regression.

**Supplementary Figure 9:**
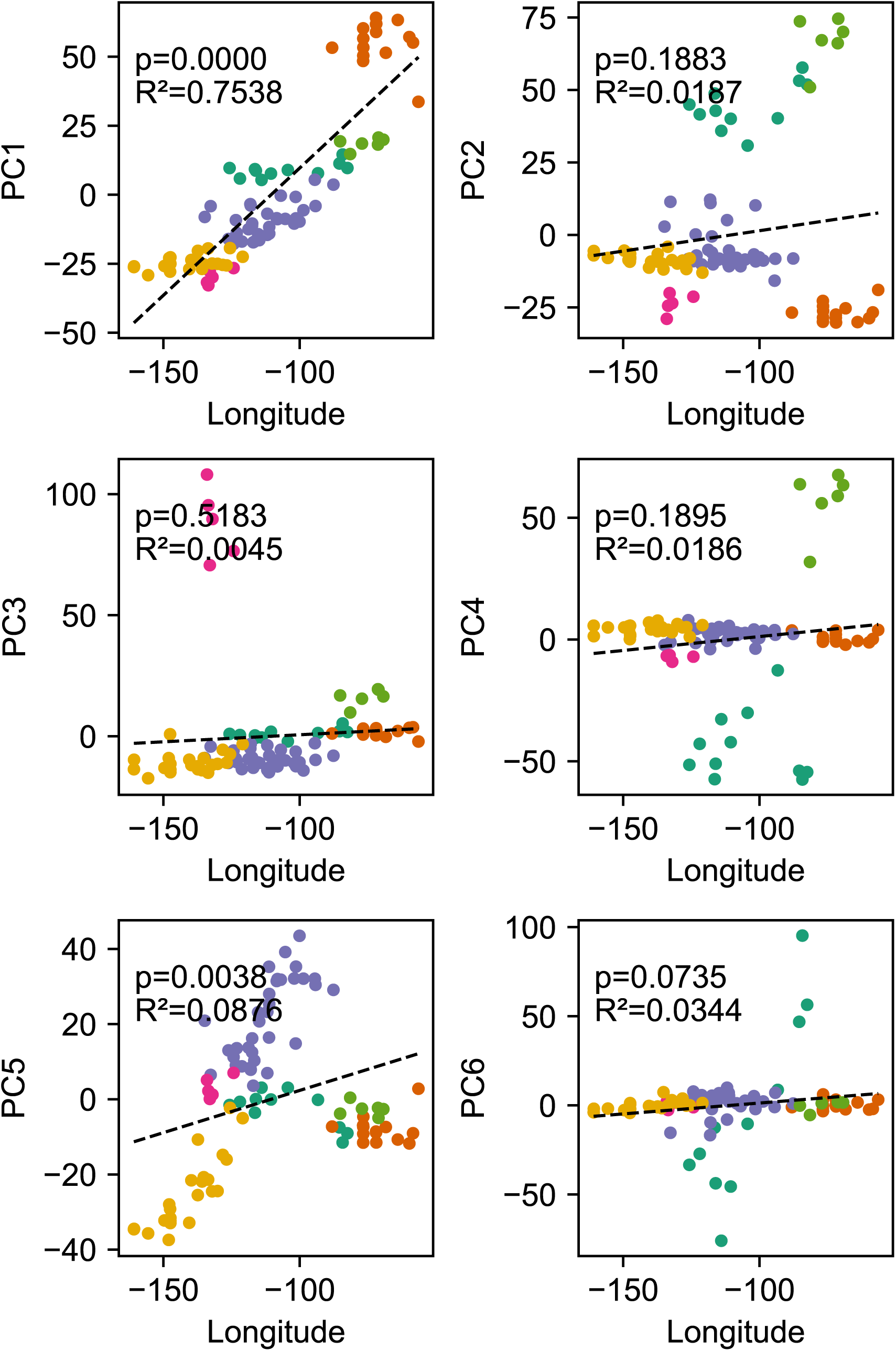
Relationship between top axes of genetic variation and longitude: In each sub-panel we plot the PC value against longitude for each sample in the gray wolf dataset. We see many of the top PCs are significantly correlated with longitude as tested by linear regression.

**Supplementary Figure 10:**
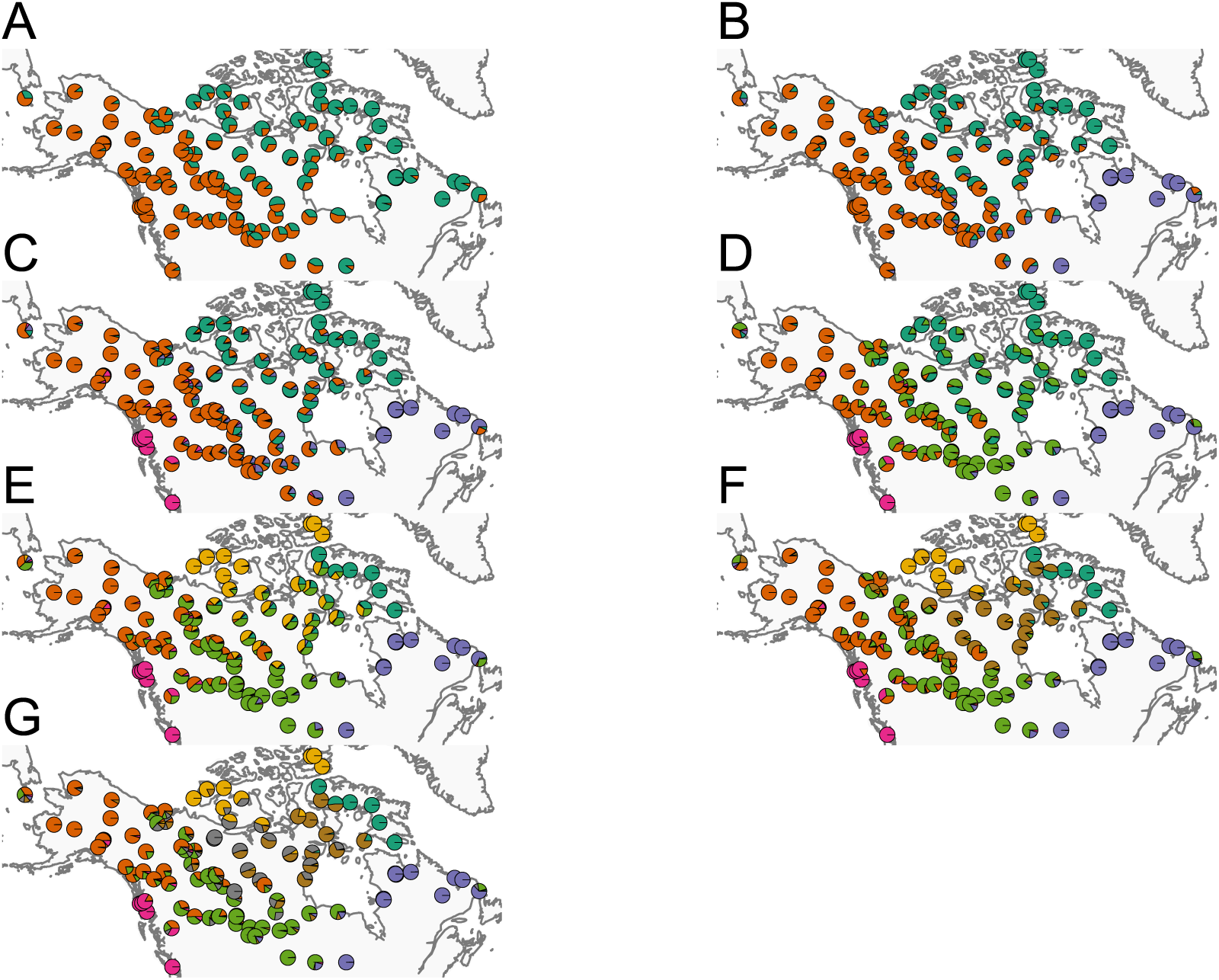
Summary of ADMIXTURE results: (A-G) Visualization of ADMIXTURE results for *K =* 2 to *K =* 8. We display admixture fractions for each sample as colored slices of the pie chart on the map. For each *K* we ran 5 replicate runs of ADMIXTURE and in this visualization we display the solution that achieves the highest likelihood amongst the replicates. The ADMIXTURE results qualitatively reveal a spatial signal in the data as admixture fractions tend to be spatially clustered.

**Supplementary Figure 11:**
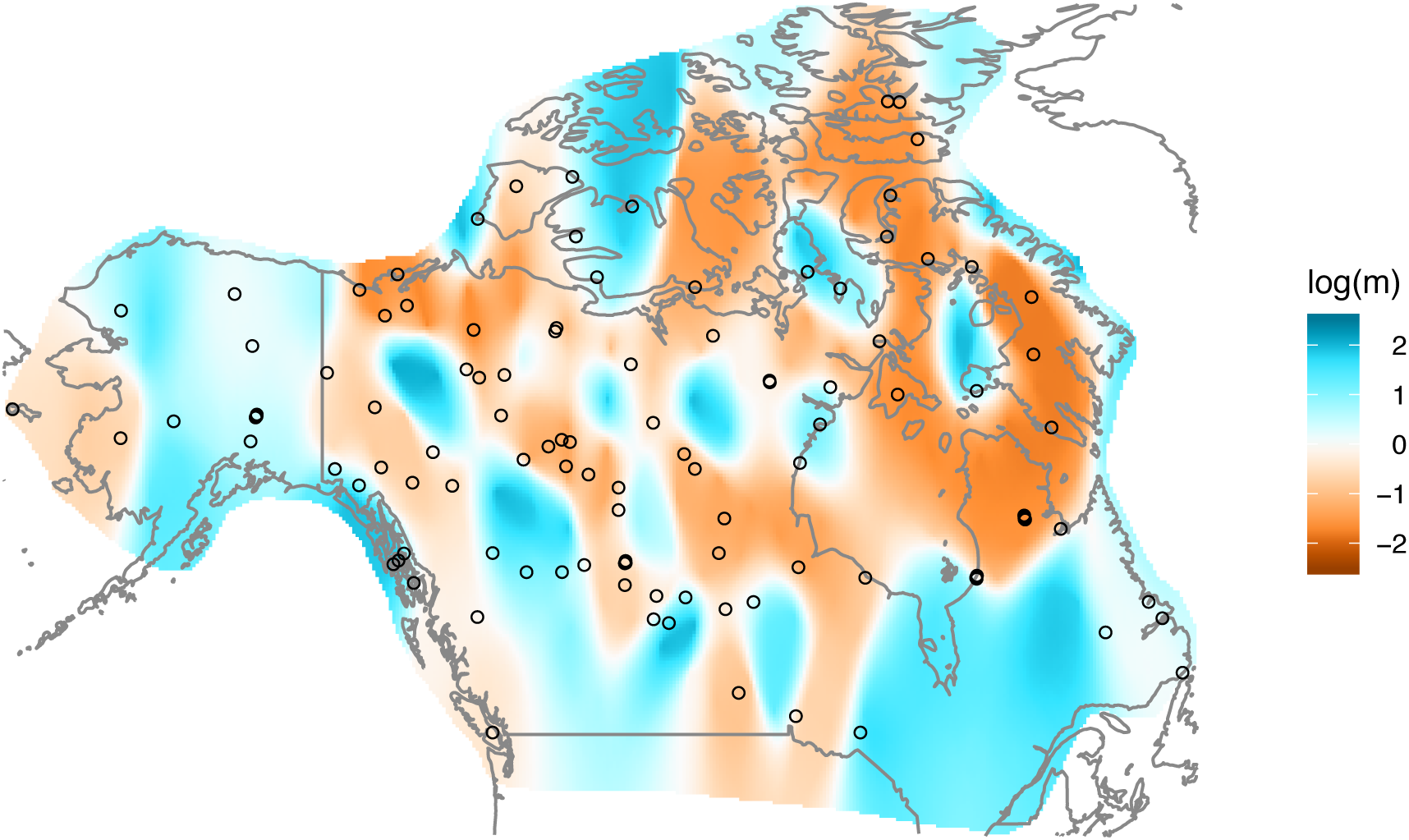
Application of EEMS to the North American gray wolf dataset: We display a visualization of EEMS applied to the North American gray wolf dataset. The more orange colors represent lower than average effective migration on the log-scale and the more blue colors represent higher than average effective migration on the log-scale. The results of EEMS are qualitatively similar to FEEMS.

**Supplementary Figure 12:**
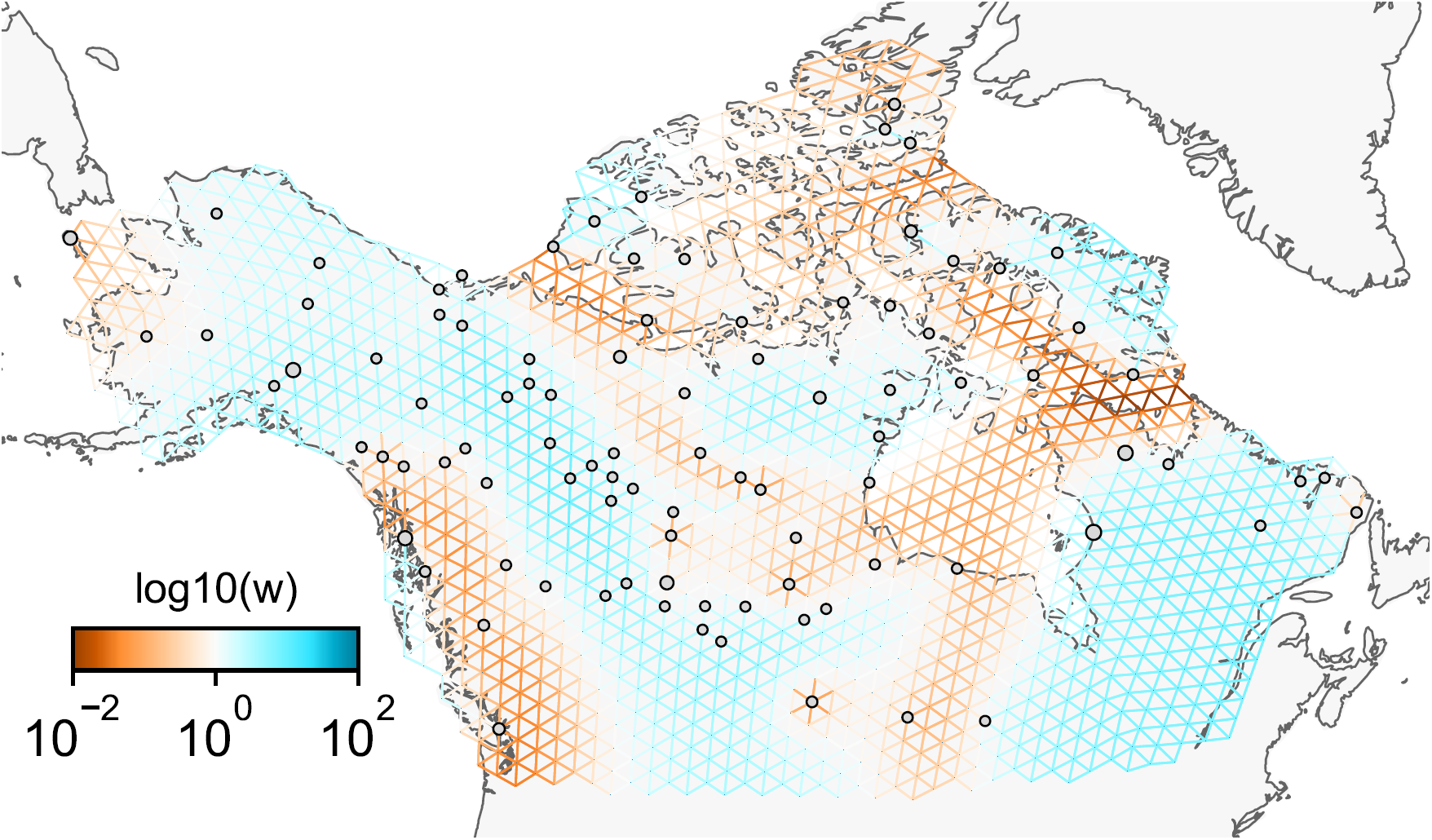
Application of FEEMS on the North American gray wolf dataset with an exact likelihood model: We display the fit of FEEMS based in the formulation (19) to the North American gray wolf dataset. This fit corresponds to a setting of tuning parameters at *λ =* 10^−3^, *α =* 50. Additionally we set the lower bound of the edge weights to ***l =*** 0.01, to ensure that the diagonal elements of ***L*** does not become too small—this has an implicit effect on 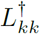, preventing it from blowing up at unobserved nodes. The more orange colors represent lower than average effective migration on the log-scale and the more blue colors represent higher than average effective migration on the log-scale. Visually the result is comparable to that of FEEMS fit (Figure 4) based in the formulation (9).

**Supplementary Figure 13:**
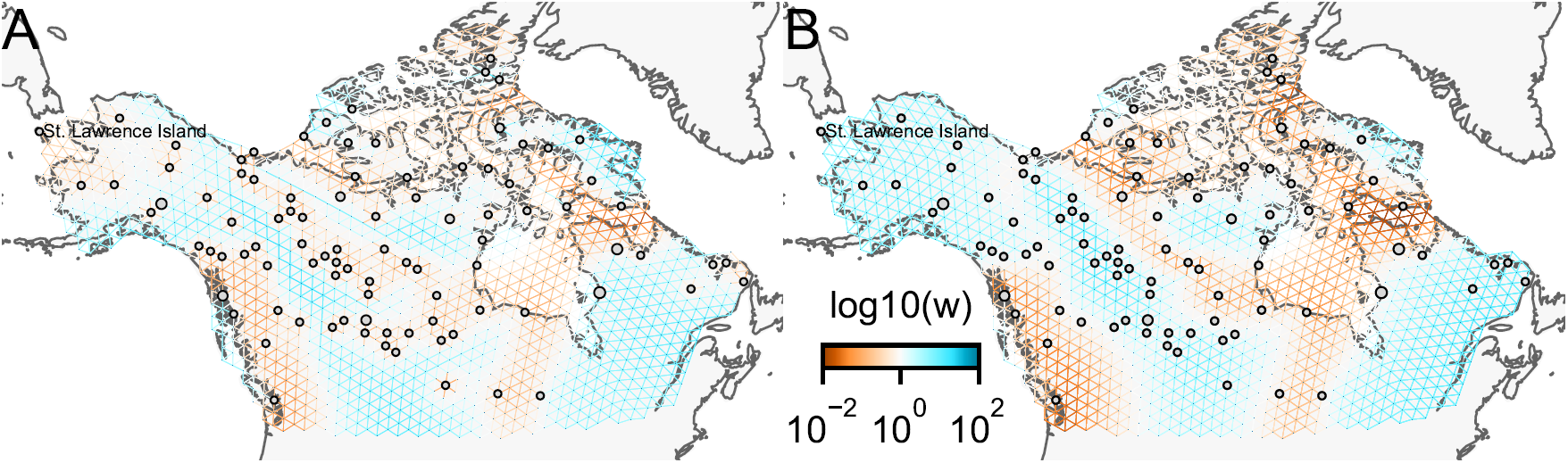
Application of FEEMS on the North American gray wolf dataset with joint estimation of the residual variance and graph’s edge weights: We show visualizations of fits of FEEMS to the North American gray wolf dataset when the residual variance and edge weights of the graph are jointly estimated. Both fits correspond to a setting of tuning parameters at *λ =* 10^−3^, *α =* 50. (A) Displays the estimated effective migration surfaces where every deme shares a single residual parameter *σ*. The result is similar to the procedure that prefixes *σ* from the homogeneous isolation by distance model (Figure 4), except the high migration edge forming long path in (A) which disappears with higher values of *α*. (B) Displays the estimated effective migration surfaces where each node has its own residual parameter *σ*_*k*_ for all nodes *k*. These node specific residual parameters allow more flexible graphs, but at the cost of over-fitting to the data. In particular, without adding smooth regularization term on the residual variances, it fails to recover some geographic features like St. Lawerence Island.

**Supplementary Figure 14:**
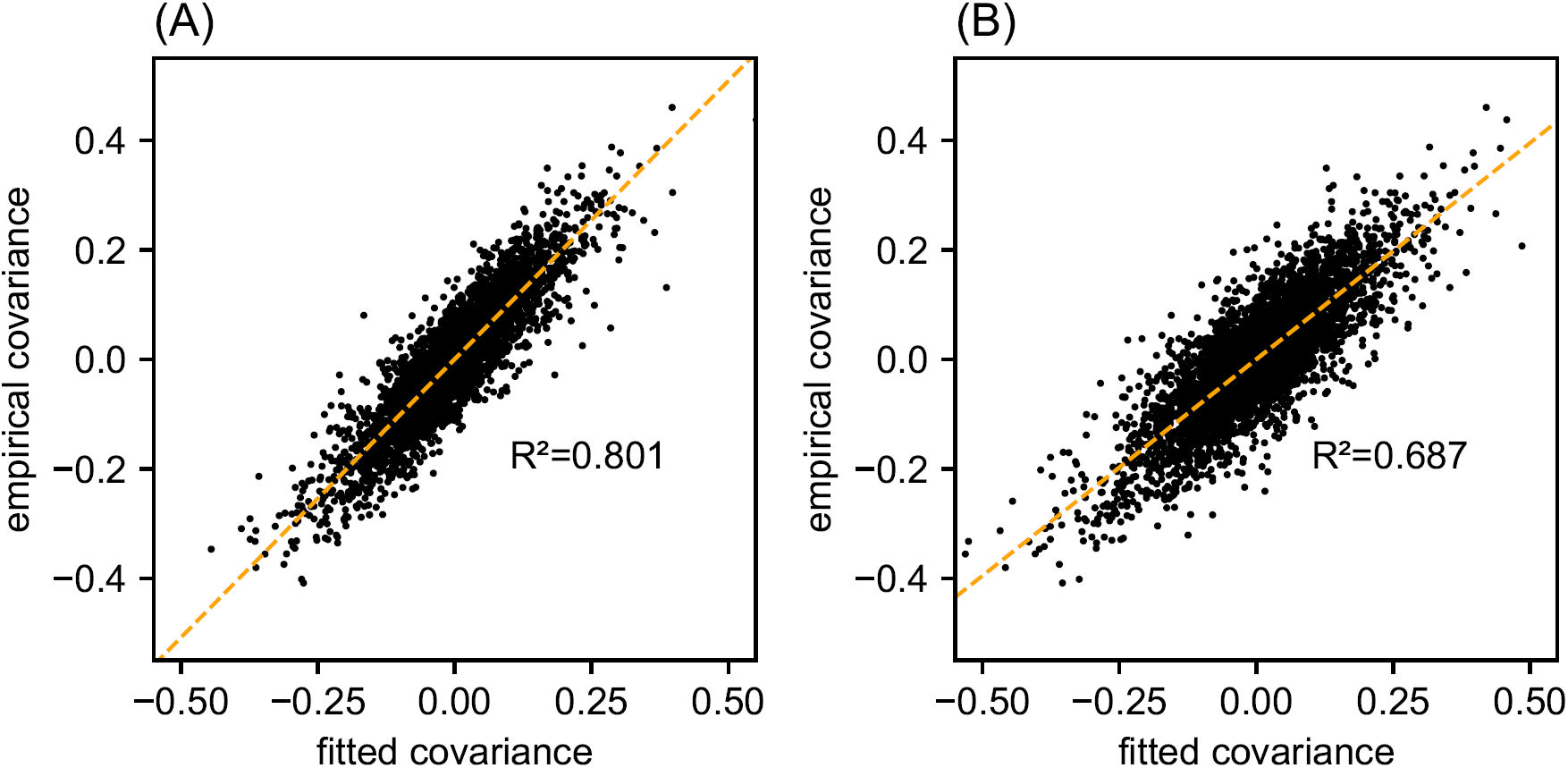
Relationship between fitted and empirical covariance on the North American gray wolf dataset: We display scatter plots of empirical genetic covariances versus fitted covariances from FEEMS fits on the gray wolf dataset. (A) Corresponds to the result shown in Figure 4. (B) Corresponds to the result shown in Supp. Fig. 13B. The x-axis represents the transformed fitted covariance matrix, i.e. 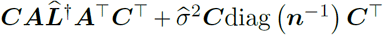 (see equation (6)). The y-axis represents the transformed sample covariance matrix, i.e.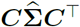. The simple linear regression fit is shown in orange dashed lines and *R*_2_ is given.

**Supplementary Figure 15:**
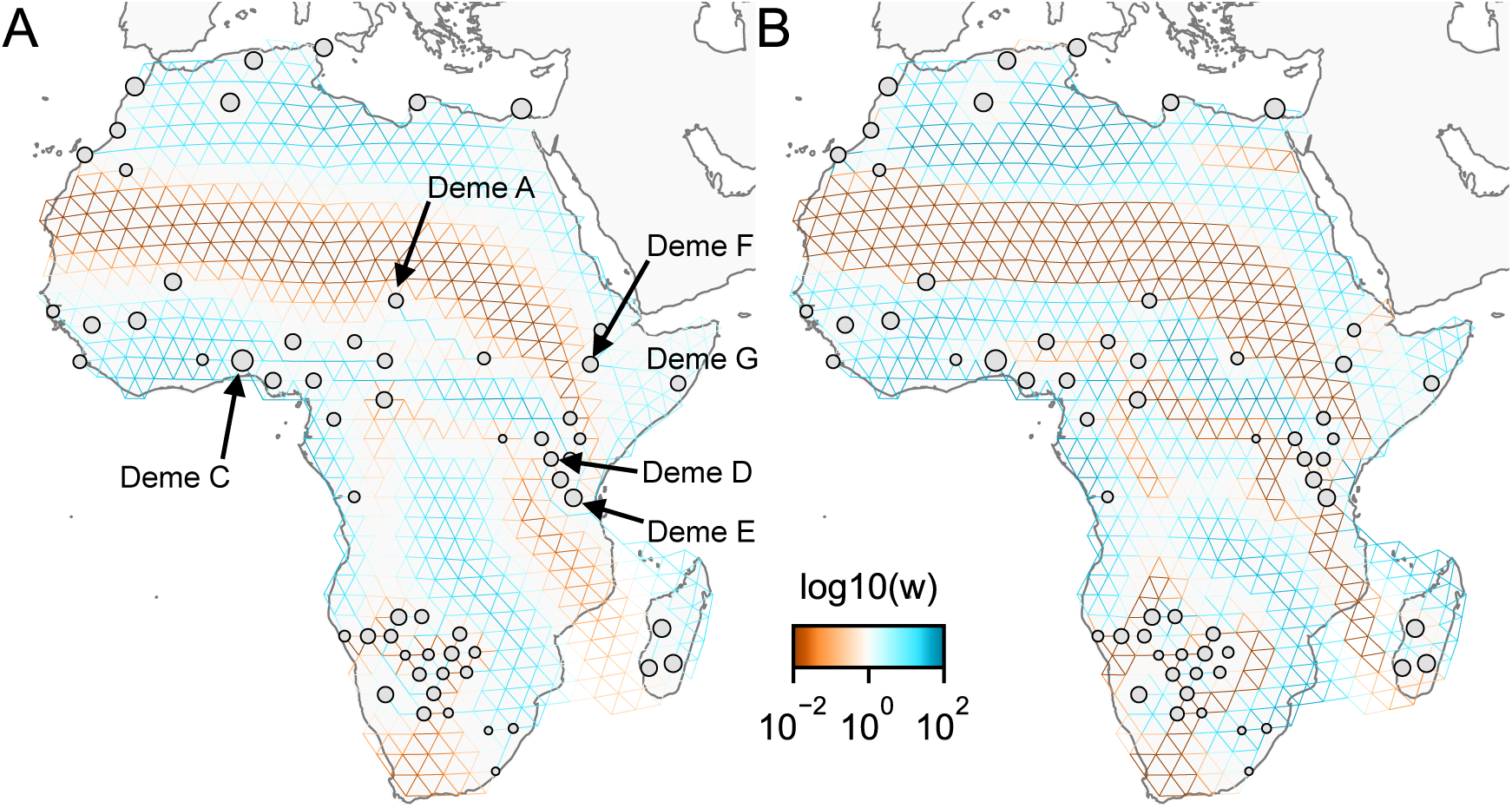
Application of FEEMS to a dataset of human genetic variation from Africa with different parameterization: We display visualizations of FEEMS to a dataset of human genetic variation from Africa with different parameterization of the graph’s edge weights. See (Peter et al., 2018) for the description of the dataset. (A) Displays the recovered graph under the edge parameterization. (B) Displays the recovered graph under the node parameterization. Both parameterization have their own regularization parameters *λ* and *α*, but these parameters are not on the same scale. We set *λ =* 2·10^−4^, *α =* 10 for the node parameterization which is seen to yield similar results to those in (Peter et al., 2018). For the edge parameterization, we keep the same *λ* value while we set *α =* 60 so that the resulting graph reveals similar geographic structure to the node parameterization. We also set the lower bound ***l*** = 0.01. From the plots, it is worth noting two important distinctions: (1) We see the migration surfaces shown in (B) recover sharper edge features while the migration surfaces in (A) are overall smoother. This is attributed to the fact that node parameterization has its own additional regularization effect on the edge weights, and in order to achieve similar degree of regularization strength for the edge parameterization, it needs a higher regularization parameters, which results in more blurring edges than the node parameterization. (2) When measuring correlation of the estimated allele frequencies among nodes, we find that Deme B is the node with the second highest correlation to Deme A, whereas Deme C (and nearby demes) is not as much correlated to Deme A compared to Deme B. Panel (A) reflects this feature by exhibiting a corridor between Deme A and Deme B and reduced gene-flow beneath that corridor. This reduced gene-flow disappears in (B), even if the regularization parameters are varied over a range of values. Additionally, Deme D is most highly correlated to Deme E, F, and G, and this is implicated by a long-range corridor connecting those demes appearing in Panel (A) while not shown in (B). These results point a conclusion that the form of the node parameterization is perhaps too strong and in this case it limits model’s ability to capture desirable geographic features that are subtle to detect.

**Supplementary Figure 16:**
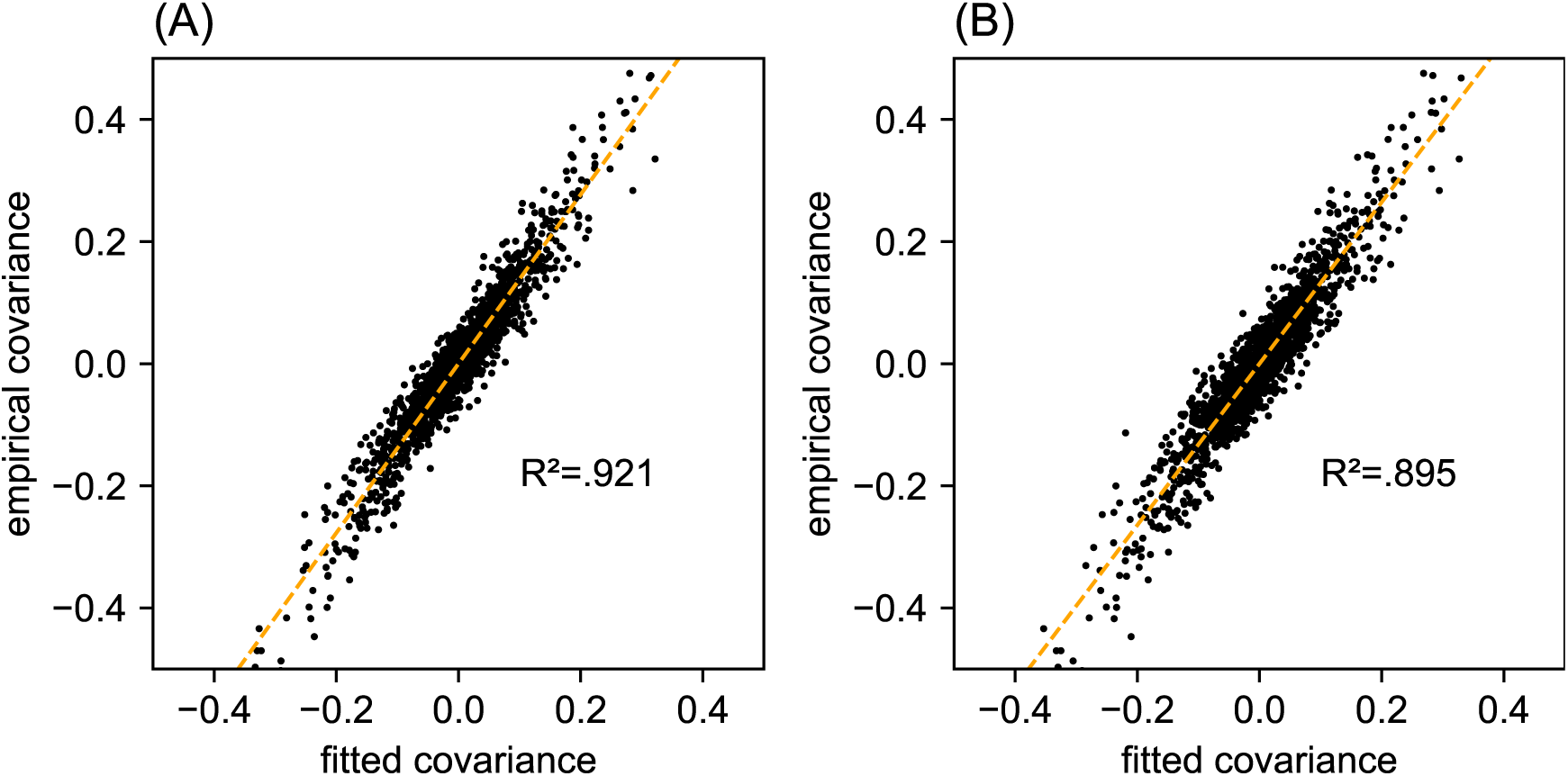
Relationship between fitted and empirical covariance on a dataset of human genetic variation from Africa: We display scatter plots of empirical genetic covariance versus fitted covariance from FEEMS fits on the African dataset. (A) Corresponds to the result shown in Supp. Fig. 15A. (B) Corresponds to the result shown in Supp. Fig. 15B. The x-axis represents the transformed fitted covariance matrix, i.e. 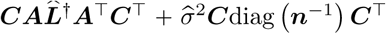 (see equation (6)). The y-axis represents the transformed sample covariance matrix, i.e.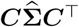. The simple linear regression fit is shown in orange dashed lines and *R*_2_ is given.

**Supplementary Figure 17:**
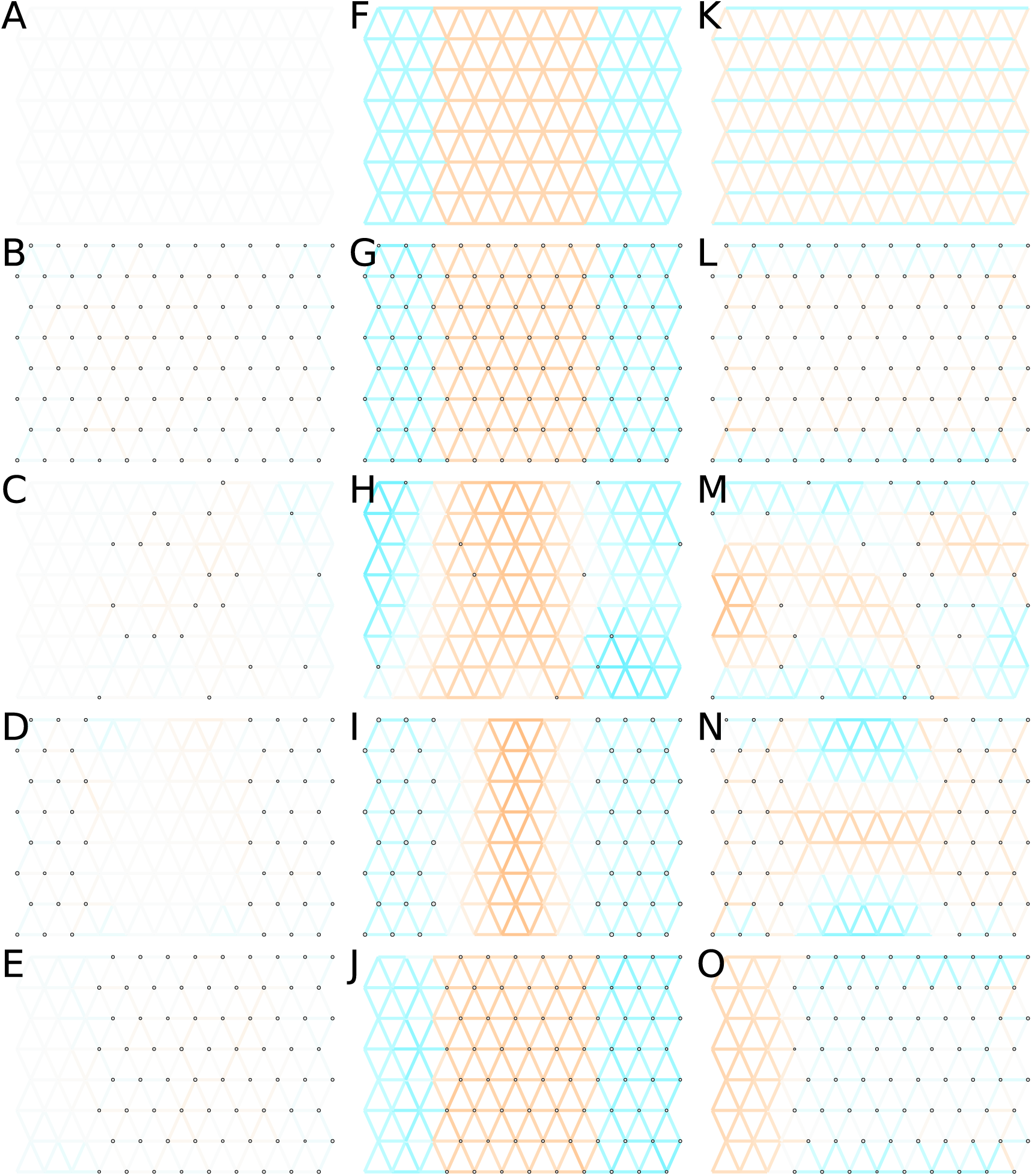
Application of FEEMS based on node parameterization to an extended set of coalescent simulations: We display an extended set of coalescent simulations with the same migration scenarios and sampling designs as Supp. Fig. 2. The sample sizes across the grid are represented by the size of the grey dots at each node. The migration rates are obtained by solving the FEEMS objective function (9) with node parameterization where the regularization parameters are specified at *λ =* 10^−3^, *α =* 50. (A, F, K) display the ground truth of the underlying migration rates. (B, G, L) Shows simulations where there is no missing data on the graph. (C, H, M) Shows simulations with sparse observations and nodes missing at random. (D, I, N) Shows simulations of biased sampling where there are no samples from the center of the simulated habitat. (E, J, O) Shows simulations of biased sampling where there are only samples on the right side of the habitat.

**Supplementary Figure 18:**
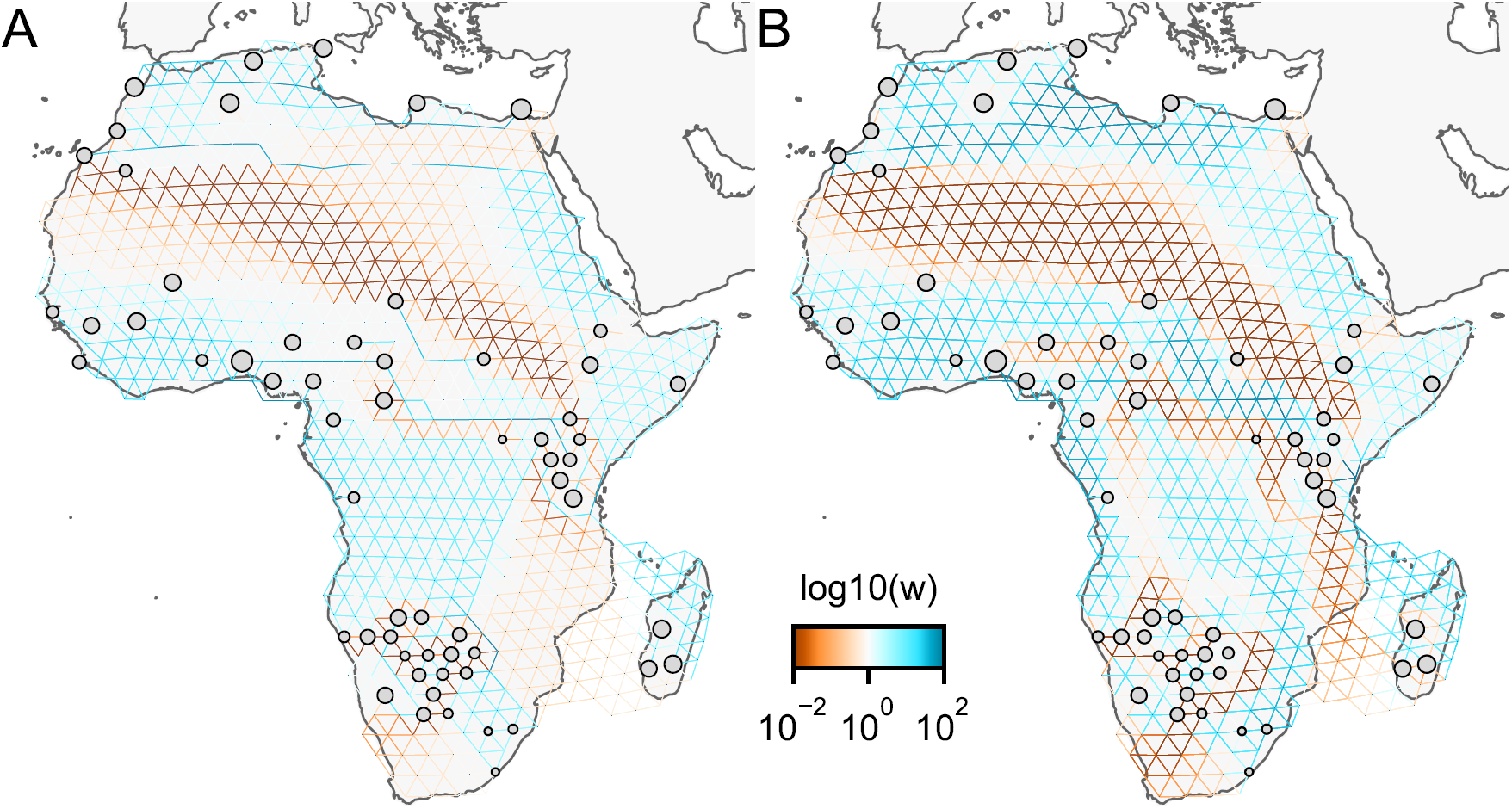
Application of ℓ_1_-norm-based FEEMS to a dataset of human genetic variation from Africa: We display visualizations of FEEMS to a dataset of human genetic variation from Africa with the ℓ_1_-based penalty function. See Peter et al. (2018) for the description of the dataset. (A) Displays the recovered graph under the edge parameterization with ℓ_1_ norm based penalty where the regularization parameters are specified at *λ* = 4·10^−2^, *α* = 30. (B) Displays the recovered graph under the node parameterization with ℓ_1_ norm based penalty where the regularization parameters are specified at *λ* = 4·10^−2^, *α =* 1. To minimize the objective (20), linearized ADMM is applied with 20, 000 number of iterations. The lower bound is set to be ***l*** = 0.01 for both parameterizations. Note that due to the high degrees of missingness, the estimated effective migration surfaces using solely ℓ_1_-based penalty exhibit many likely artifacts (e.g., high migration edges forming long paths, seen in A) unless an additional penalty term is added to promote global smoothness of the edge weights such as a combination of ℓ_1_ norm penalty function and node parameterization as shown in (B).

**Supplementary Figure 19:**
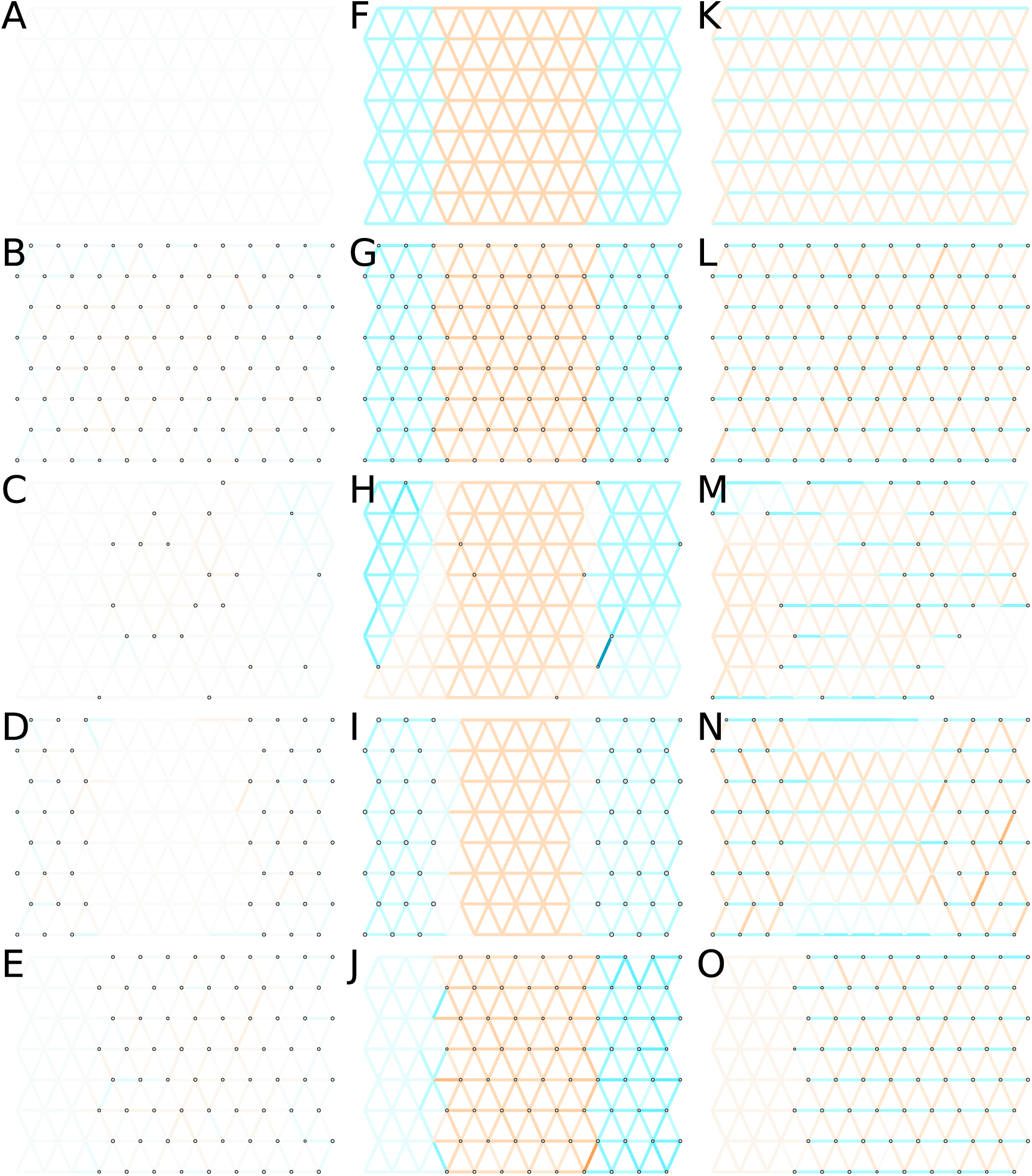
Application of ℓ_1_-norm-based FEEMS to an extended set of coalescent simulations: We display an extended set of coalescent simulations with the same migration scenarios and sampling designs as Supp. Fig. 2. The sample sizes across the grid are represented by the size of the grey dots at each node. The migration rates are obtained by solving ℓ_1_ norm based FEEMS objective (20) where the regularization parameters are specified at *λ =* 10^−1^, *α =* 30 (I), *λ =* 10^−3^, *α =* 30 (N), and *λ =* 10^−2^, *α =* 30 for the rest. (A, F, K) display the ground truth of the underlying migration rates. (B, G, L) Shows simulations where there is no missing data on the graph. (C, H, M) Shows simulations with sparse observations and nodes missing at random. (D, I, N) Shows simulations of biased sampling where there are no samples from the center of the simulated habitat. (E, J, O) Shows simulations of biased sampling where there are only samples on the right side of the habitat.

## Footnotes

1 Specifically, since we use a triangular grid embedded in geographic space to define the graph 𝒢, the pattern of nonzero elements is prefixed by the structure of the sparse traingular grid.

2 To be more precise, under (2), (3), the law of total variance formula leads to specific formulas for the mean and variance structure as given in (4), whereas the marginal distribution of 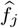 is not necessarily a Gaussian distribution. We simply chose the Gaussian distribution here to enable easy calculation for the data likelihood. We believe the specific choice of the likelihood is not that critical as long as the first two moments of the distribution can be matched closely.

3 Our model (6) says that the *p* SNPs are independent. This assumption is unlikely to hold when SNPs are in close chromosomal proximity are analyzed due to linkage disequilibrium. In (Petkova et al., 2016), they introduce the effective degree of freedom *ν* ∈[*o−*1, *p*] to account for such dependency and instead consider the model 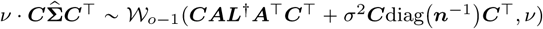 with *ν* being estimated alongside other model parameters. In FEEMS, we note that the degree of freedom parameter does not affect the point estimate produced by our algorithm.

4 We remark that besides the effective degree of freedom and the SNP-specific re-scaling by *µ*_*j*_ (1 −*µ*_*j*_), the EEMS (Petkova et al., 2016) and FEEMS likelihoods are equivalent up to constant factors, as long as only one individual is observed per node and the residual variance *σ*^2^ is allowed to vary across nodes—See Supp. Note “ *Jointly estimating the residual variance and edge weights* “ for details. In addition, constant factors are effectively absorbed into the unknown model parameters ***L*** and *σ*^2^ and therefore it does not affect the estimation of effective migration rates, up to constant factors.

5 We solve using linearized ADMM when the penalty function is l_1_ norm, i.e. *λ ∥***Δ ((***w) + α* log ((*w))∥*_1_ (Boyd et al., 2011).

6 More precisely, it is possible to achieve 𝒪(d*o + o*^3^) per-iteration complexity if one employs a solver that is specially designed for sparse Laplacian system. In our work we use sparse Cholesky factorization which may slightly slow down the per-iteration complexity. See Supp. Material for the details of the gradient and objective computation.

